# Deep learning methods in metagenomics: a review

**DOI:** 10.1101/2023.08.06.552187

**Authors:** Gaspar Roy, Edi Prifti, Eugeni Belda, Jean-Daniel Zucker

## Abstract

The ever-decreasing cost of sequencing and the growing potential applications of metagenomics have led to an unprecedented surge in data generation. One of the most prevalent applications of metagenomics is the study of microbial environments, such as the human gut. The gut microbiome plays a crucial role in human health, providing vital information for patient diagnosis and prognosis. However, analyzing metagenomic data remains challenging due to several factors, including reference catalogs, sparsity, and compositionality. Deep learning (DL) enables novel and promising approaches that complement state-of-the-art microbiome pipelines. DL-based methods can address almost all aspects of microbiome analysis, including novel pathogen detection, sequence classification, patient stratification, and disease prediction. Beyond generating predictive models, a key aspect of these methods is also their interpretability. This article reviews deep learning approaches in metagenomics, including convolutional networks (CNNs), autoencoders, and attention-based models. These methods aggregate contextualized data and pave the way for improved patient care and a better understanding of the microbiome’s key role in our health.

**Author summary:** In our study, we look at the vast world of research in metagenomics, the study of genetic material from environmental samples, spurred by the increasing affordability of sequencing technologies. Our particular focus is the human gut microbiome, an environment teeming with microscopic life forms that plays a central role in our health and well-being. However, navigating through the vast amounts of data generated is not an easy task. Traditional methods hit roadblocks due to the unique nature of metagenomic data. That’s where deep learning (DL), a today well known branch of artificial intelligence, comes in. DL-based techniques complement existing methods and open up new avenues in microbiome research. They’re capable of tackling a wide range of tasks, from identifying unknown pathogens to predicting disease based on a patient’s unique microbiome. In our article, we provide a very comprehensive review of different DL strategies for metagenomics, including convolutional networks, autoencoders, and attention-based models. We are convinced that these techniques significantly enhance the field of metagenomic analysis in its entirety, paving the way for more accurate data analysis and, ultimately, better patient care. The PRISMA augmented diagram of our review is illustrated in **Fig 1**.

## Introduction

The human body hosts a vast number of different microorganisms species (bacteria, viruses, archaea, fungi and protists) who dwell inside and on our bodies according to complex interactions, not only among each other but also with their host, forming complex ecosystems. This entire habitat, including the microorganisms, their genomes and the surrounding environment is called the “microbiome”, while the whole genetic material is referred to as the “metagenome” ([1]). The gut microbiome for instance, plays a key role in the functioning of our own organism and is considered a “super-integrator” of patient health ([2]). The lack of microbial diversity is an indicator of chronic disease of the host (see [3], [4] and [5]), but also of the health evolution after an intervention (see [6] and [7]). It is therefore important to develop tools that allow to characterize and understand both its composition and its links with human health and disease.

The common approach used to explore the microbiome is to start by charting the species that compose it, quantify their diversity as well as their abundance and eventually, their functional potential. This is made possible by advances of Next-Generation Sequencing (NGS) technologies, without the need to cultivate specific organisms. These technologies have opened the way to the genetic characterization of entire ecosystems and have accelerated the now rapidly growing field of metagenomics. Typical metagenomic data consist of millions of reads that can then be associated to the microorganism they originate from using reference catalogs.

In order to assert the species composition of a metagenome, two main approaches (shown in the first step of **Fig 2**) are widely used to characterize microbial communities with high-throughput sequencing, producing DNA reads of microbes. However, different studies typically use different regions, making it difficult to compare results between studies.

**Fig 1.**
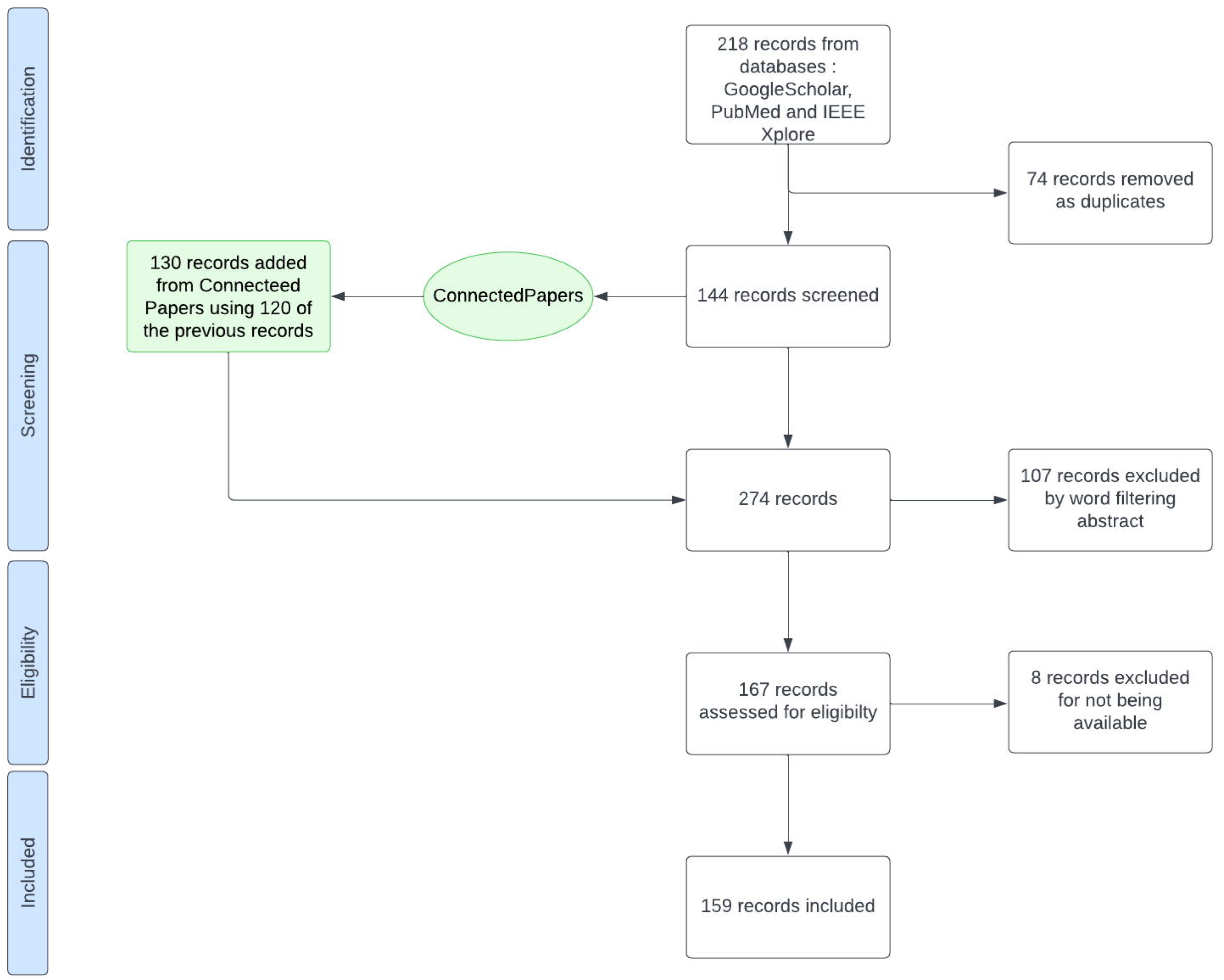
PRISMA-type diagram for article selection of this review. The method developed here enriches the research equation selection with Connected Papers, this diagram represents the selection along with this enrichment in green

**Fig 2.**
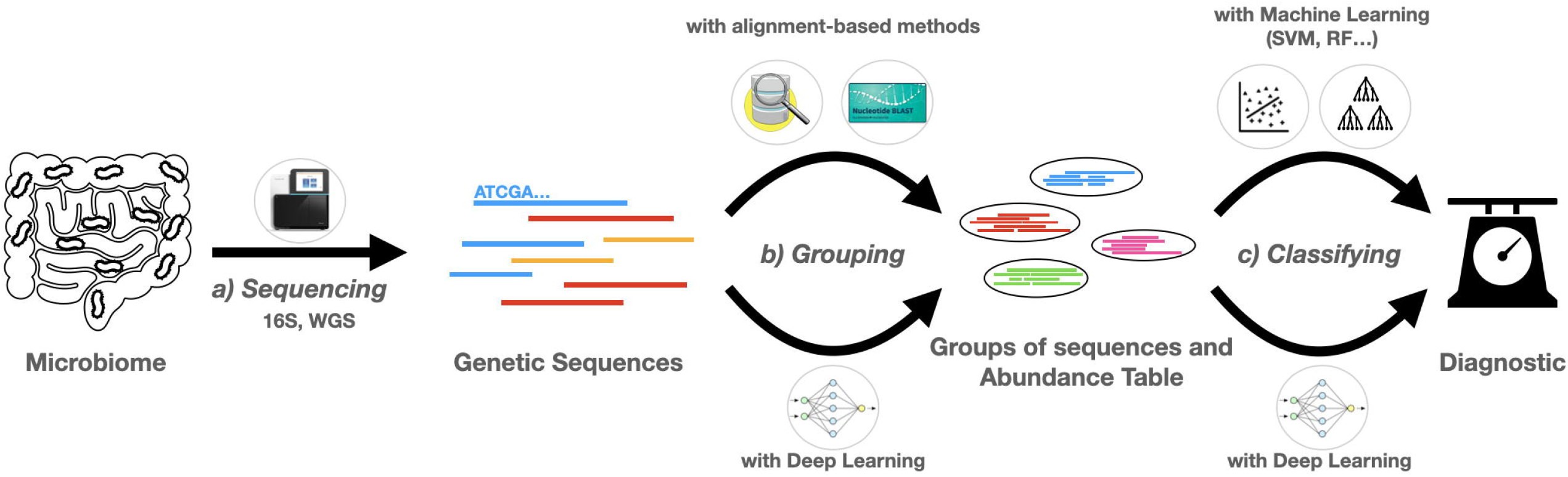
Illustration of the use of deep learning in disease prediction from metagenomic data. The classic simplified pipeline for disease prediction from microbiome data follows three distinct steps. In step a), high-throughput sequencing of DNA libraries from environmental samples generates millions of reads (from whole genomic DNA in WGS metagenomics or from targeted 16S rRNA genes in targeted metagenomics) from the organisms that make up the community. Second, in step b), the sequences are either clustered or classified into different groups to characterize the different species present in the sample. This step can be realized by classical bioinformatics pipelines, such as alignment-based methods, or by more recent DL architectures, both of which can be used to estimate their relative abundance. In step c), the abundance table or the embeddings extracted from the use of NNs can be used to classify the metagenomes as coming from patients with the disease state or not. DL methods can also be used to integrate additional information (annotations, genes, phylogeny, etc.) to classify sequences or metagenome profiles.

Sequencing a relatively short DNA region requires a low number of reads, resulting in inexpensive analyses. 16S sequencing has been pivotal in the characterization of microbial ecosystems and is still widely used in quantitative metagenomic studies, despite known drawbacks associated to the variability in diversity estimates and taxonomic resolution of different hypervariable regions ([8], [9]), the lack of resolution at lower taxonomic levels than genus and the fact that functional information about the ecosystem can only be indirectly inferred ([10]). Efforts in sequencing full-length 16S genes with third-generation sequencing technologies shows better taxonomic resolution ([11]). However, 16S rRNA is completely useless for viral data analysis methods because viruses do not have any of these genes.

The second method is Whole Genome Shotgun (WGS) metagenomics, which sequences and analyses the entire genomic content of all organisms in the environmental sample. This makes it possible to characterize the full diversity of the ecosystem, including archaea, bacteria, viruses, and eukaryotes. Unlike 16S, WGS data are highly resolutive and more complex, enabling differentiation down to the strain level as well as direct functional potential profiling. ([12] and [13]). However, the short read technologies used can make it challenging for the bioinformatics pipelines to classify sequences. It is expected that future sequencing technologies will further increase the popularity of the WGS approach in microbiome studies.

Both methods are still widely used, although WGS is gaining popularity as sequencing costs decreases and the development of more powerful and faster bioinformatics pipelines are developed to extract more knowledge from the data.

Sequencing can produce several forms of reads, including raw reads (as sequenced) and contigs. Contigs are longer sequences generated by assembly tools and are often used to discover genes, that make up more complex reference catalogs. Sequenced reads can be short (less than 300 bp) or long (today with an average between 10 to 30k bp)([14]) - both having their advantages and drawbacks.

It is important here to remember the concept of k-mer. A k-mer is a subsequence of a DNA sequence of length k. In most cases, k-mers in a sequence are considered to be overlapping, i.e. the first k-mer consists of the k first bases of the sequence, while the second k-mer consists of the bases from the second to the k+1th and so on. K-mers are important when looking at a DNA sequence because they can capture important patterns of the molecule.

All of these sequences are then analyzed to achieve different goals. A first goal may be to identify sequences of interest such as those associated with specific functions. This task will be referred to as “functional annotation”. Some methods involve processing each read individually to search for specific sequences associated with pathogens or other global functions.

However, a key issue in metagenomics is also to identify which microorganisms are actually present in the sample. This can be achieved by performing either de-novo metagenomic assembly of metagenomic reads or assembly-free approaches where metagenomic reads are used directly for taxonomic and functional profiling based on reference databases.

To identify and reconstruct the genomes of the species that make up the microbiome, raw reads are first assembled into *contigs*, which carry more information. However, their assembly is prone to error because overlapping sequences may be slightly different, requiring a consensus sequence. The sequences are then grouped, or “binned”, either in a supervised manner using alignment to genomes of reference for example ([15]), or in an unsupervised manner, independent of reference sequences, exploiting other sources of information like compositional profiles such as k-mer distribution and abundance profiles ([16],([17]), [18] and [19]). By binning contigs, it is possible to reconstruct whole or part of the genome of some of species present in the metagenome. The resulting genome is called a Metagenome-Assembled Genome (MAG) of Metagenomic Species (MGS) ([19])). In this context, the human gut microbiome is one of the microbial ecosystems that has been more extensively characterized at the genomic level, with several large-scale metagenomic assembly studies yielding comprehensive catalogs of human gut MAGs ([20], [21], [22]). When using MAGs, it is also possible to calculate the relative abundance of each MAG in the metagenome by considering the number of reads mapped to a MAG. In both cases, this results in an abundance table representing the metagenome by the abundance of each species. Another approach is to start by building representative, non-redundant gene catalogs ([23] [24]), which are themselves binned to Metagenomic Species (MGS)([19]) and ([25])). At the end of this step, the output is an abundance table linking each taxon to its Metagenomic Gene Abundance (MGA).

Other methods, called “assembly-free methods” start by grouping together the reads that belong to a particular taxonomic unit, such as species. They exploit sequence similarity ([15], [26], [27]) or kmer-content similarity ([16], [28]) against reference databases. For example, reads are aligned against gene markers for taxonomic profiling ([29]) or comprehensive gene catalogs that maximize the genomic knowledge of microbial ecosystems, such as Genome Taxonomy Database (GTDB), the Global Microbial Gene Catalog (GMGC) or the Kyoto Encyclopedia of Genes and Genomes (KEGG). This provides a representation of the composition of a metagenome as well as its functional potential. The number of reads in each bin provides an estimate of their relative abundance within the metagenome, when normalized by the respective size of their respective genomes.

Traditional bioinformatics methods, while very useful and widely used, have several drawbacks: they can be computationally expensive, are affected by sequencing errors and are often dependent on reference databases. However, the majority of the microorganisms found in the human microbiome remain poorly characterized. Over the last decade, new methods have been developed in the field of sequence classification that have enabled several breakthroughs. SVM or Random Forest based methods have proven their efficiency and are now very good alternatives to alignment based methods in order to classify sequences ([30])

Although the primary goal of both contig-based and assembly-free methods remains the reconstruction of the metagenome at the species level, they can also be used to obtain an abundance table representing the distribution of the species that make up the metagenome. This way of handling reads to obtain a quantification of the microbiome can be referred to as “quantitative methods”. Finally, once the abundance table of the metagenome is obtained, it can be used for disease prediction analyses. More specifically, this consists of establishing links between the metagenomic data obtained in the first step and patient information such as disease status or severity. A brief summary of these steps is illustrated in **Fig 2**.

The task of predicting patient phenotype can be addressed using various Machine Learning (ML) models. With an increasing number of public example datasets, these algorithms can learn and extract important patterns from the data in order to classify samples based on their various characteristics. Deep Learning (DL) is a specific branch of ML that focuses on algorithms based on layers of artificial neurons that receive and process information from previous layers of neurons ([31]). The creation of diverse network types hinges on the choice of layers, neurons, and their respective organization. These range from traditional multi-layer perceptrons to recurrent neural networks, which excel at managing data that evolves over time. ([32]). Data is channeled through the network to generate an output, facilitating the learning process as the network adjusts the neuron weights via backpropagation of errors. The most notable strides empowered by deep learning are discernible in domains like image recognition and Natural Language Processing.(NLP).

Deep Learning stands out for its superior performance on large datasets, outdoing many other machine learning algorithms that reach a performance plateau with a given quantity of data. Furthermore, deep learning techniques possess a robust capacity to unearth intricate features, often imperceptible to human observation. This concept, known as “representation learning,” involves automatic discovery of necessary representations for feature detection or classification directly from unprocessed data. DL can also perform various learning paradigms (unsupervised ([33]), semi-supervised ([34]), Multiple Instance Learning ([35])), etc. These paradigms allow different types of learning: exploring the data in a certain direction with supervised learning, letting the network the task to draw conclusions with unsupervised learning, etc. In particular, the ability to learn mathematical representations from the data, such as numerical vectors called “embeddings”, makes it possible to group or mathematically classify different samples or observations. An embedding is a low-dimensional vector representation of high-dimensional data, such as sequences in genomics, which capture semantic and syntactic relationships between the elements being embedded. They are used to translate high-dimension data that would be difficult to work with for a ML model. There are different ways to embed data, especially metagenomic data, from vectors extracted with attention to representing metagenome abundance data as images in order to analyse it ([36]. They can then be used for clustering or classification. In the field of Natural Language Processing (NLP), the dimensions of typical embeddings can vary from 50 to 300. The popular Word2Vec embeddings are available in sizes of 50, 100, 200, and 300 dimensions. However, in more complex models like the original Transformer, the embedding size is 512. For even more advanced models like GPT-3, the largest variant can have an embedding size as large as 12288. Therefore, the term “low” is relative and depends on the context.

Various types of Neural Networks (NN) find extensive application in metagenomics. Notably, we can reference the conventional feed-forward Neural Network, also known as the Multi-Layer Perceptron ([37]). In this model, data flow is unidirectional, with each layer comprising a specific number of neurons interconnected to all neurons in the preceding and succeeding layers. While this straightforward form of neural network demonstrates remarkable results in data classification, it can encounter challenges tied to the data’s structure, such as overfitting, vanishing gradient problems, local minima, etc.

Convolutional Neural Networks (CNN) ([38]) are well known for their performance in image classification. Inspired by the cortex of vertebrates, they use the operation of convolution to extract spatial features. In the case of metagenomics, they can be used to classify sequences with common local patterns, such as common nucleotide patterns, but also to characterize the global structure of the microbiome.

Furthermore, Recurrent Neural Networks (RNN) ([39]), with the introduction of cycles in connections, are well suited for temporal or sequential data processing. Today, the most widely used version of RNN is the Long Short-Term Memory Neural Network (LSTM), which performs better at detecting long-term dependencies. For example, these networks can be employed to analyze DNA sequences, enabling predictions about the presence of specific DNA elements, like phages ([40]). Or they can be used to analyze the abundance of microbial species through time to predict for instance the evolution of the microbial ecosystem ([41] and [42]).

Autoencoders are a type of neural network designed to distill pertinent features from input data [43]. Their operation involves dimensionality reduction of the input data (encoding) followed by its reconstruction from the encoded data (decoding). If the decoding process proves efficient, the encoded features are extracted, offering a fresh representation. It is interesting to use this new representation in order to simplify the data and make it suitable for classification by ML algorithms or Multi-Layer Perceptron, but also to underline important features characterising the data that would not be easy to uncover otherwise. There are many types of autoencoders using various processes (variational [44], convolutional [45]…). These different types of autoencoders explore the data differently, granting access to different representations.

Another field where DL has shown remarkable results is Natural Language Processing (NLP), focused on the interactions between human and computers using natural language. Researchers have explored ways to represent, understand, analyze and generate language with AI. The biggest advances have come with the use of Transformers ([46]), a type of DL model that relies on attention mechanism to find coherence between different parts of data, one of the most famous applications being to encode the data contained in a sentence through the relations between its words (or other script sub-units).

A primary challenge in Deep Learning is the need for substantial volumes of data to train models. Given that these models comprise millions to billions of neurons, they necessitate a large number of examples to autonomously discern abstract features. In addition to procuring costly medical data, several strategies are adopted such as data augmentation or data generation methods, some of which leverage Deep Learning techniques themselves.

A critical challenge in the medical domain is not only establishing a diagnosis, but also comprehending the rationale behind it. This understanding aids in contrasting the diagnosis with a practitioner’s personal knowledge and bolsters their confidence in the outcomes. The ‘black box’ characteristic of Deep Learning models presents an obstacle here. The complexity of these models obscures the logic driving their decision-making process, underlining the significance of ‘interpretability’ in the field of Deep Learning [47]. In classic ML for example, interpretability aims not only to provide a diagnosis, but also to discover the importance of each microbiome feature in the decision process. Methods like *Predomics* [48] allow discovering highly predictive and very simple models that generalize well, while providing clear focus on the importance of the features involved. Some interesting reviews of these methods have been done by [49] and [50].

Finally, in the specific context of metagenomics, ML faces different problems, including the high-dimensional nature of the data compared to the number of samples, the vast sparsity in the data and their compositionality nature. We will explore strategies to address these challenges, optimizing the manner in which neural networks utilize the data.

In this review, we will present different DL methods used in metagenomics and analyze their motivation, qualities and drawbacks. This study focuses on the task of disease prediction itself, which is closely related to the issues of sequence classification (binning, taxonomy, identification) and ultimately phenotype prediction. Therefore, our work covers all steps and tasks performed for the analysis of human metagenome in this context. Although various reviews on Deep Learning in metagenomics exist, none of them studies all methods from raw reads to disease prediction, and they either include shallow Machine Learning and do not focus on DL, and focus on a specific metagenomic task (phenotype prediction, binning or sequence classification…).

## Materials and methods

Both the metagenomics and DL fields are currently very active, with an abundance of literature that is often not easy to navigate. We believe that this review is needed to help the reader chart the major advances while allowing for reproducible results. Our selection aimed to follow strict and reproducible rules. The pipeline of our review selection is described in **Fig 4**

### Review search equation

The first step (step (A) in **Fig 4**) of the review approach employed here consists of searching articles in three different bibliometric databases (Google Scholar, PubMed, and IEEE Xplore). This research includes the latest research until July 2023. The research equation was the following:

> Allintitle: (metagenome OR metagenomics OR metagenomic OR microbiome) AND (“deep learning” OR “neural network” OR embedding OR interpretable OR autoencoders OR CNN OR convolutional OR LSTM OR “long short-term memory” OR NLP OR “Natural Language Processing” OR transformer OR BERT)

To ensure that the requested papers cover both metagenomics and deep learning concepts, we have split our equation into two explicit parts, connected by a conjunction, focusing on metagenomics and deep learning, respectively. We decided to make different DL methods explicit because some articles directly mention the specific models they use, such as CNNs or LSTMs. This equation is summarized by **Fig 3**.

**Fig 3.**
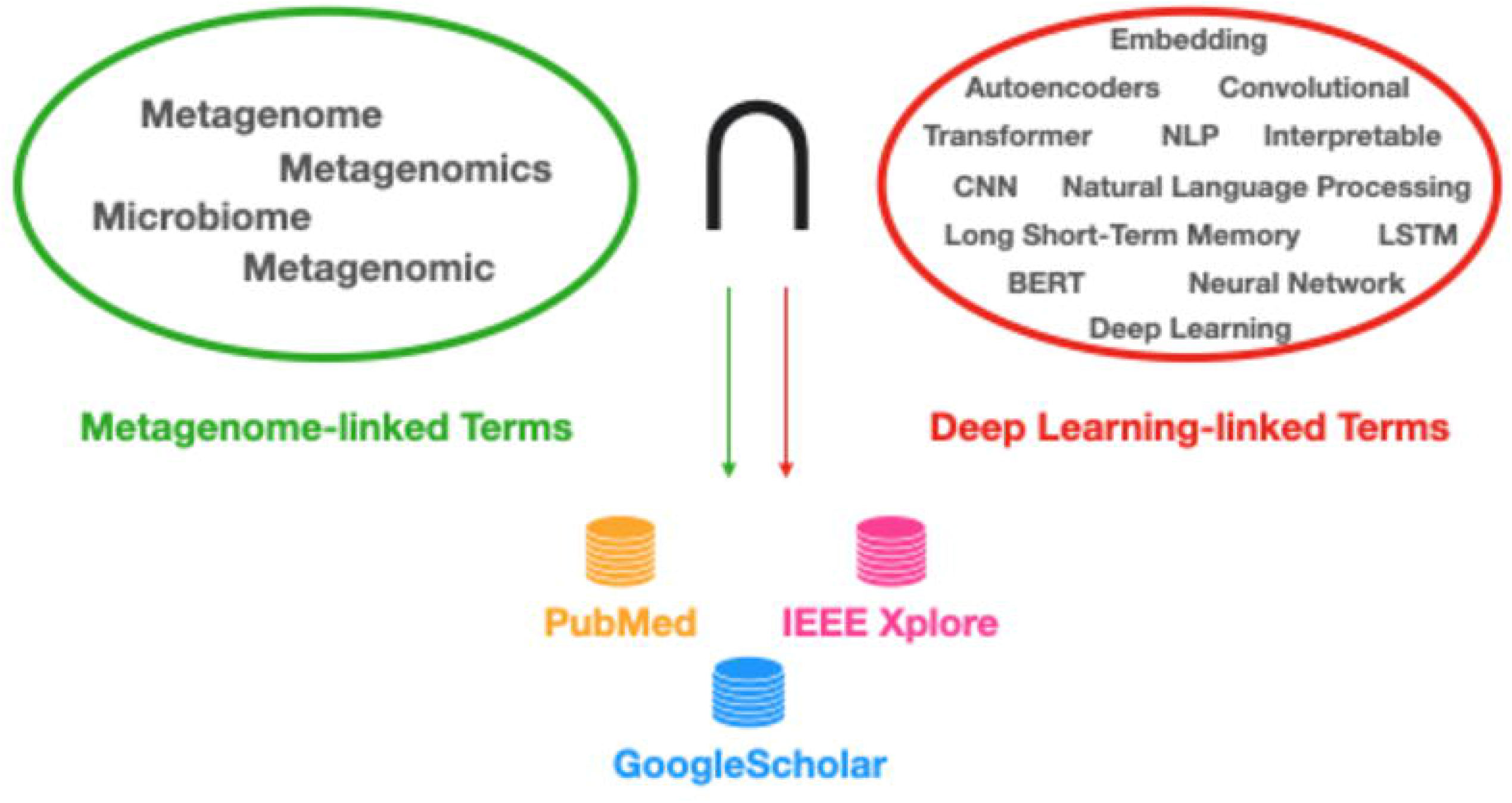
Diagram representing the structure of the research equation. Each ellipse represents a part of the research equation, respectively the metagenomics (colored in green) and DL concepts (colored in red. The equation was applied to three databases: PubMed, IEEE Xplore and Google Scholar were querried.

**Fig 4.**
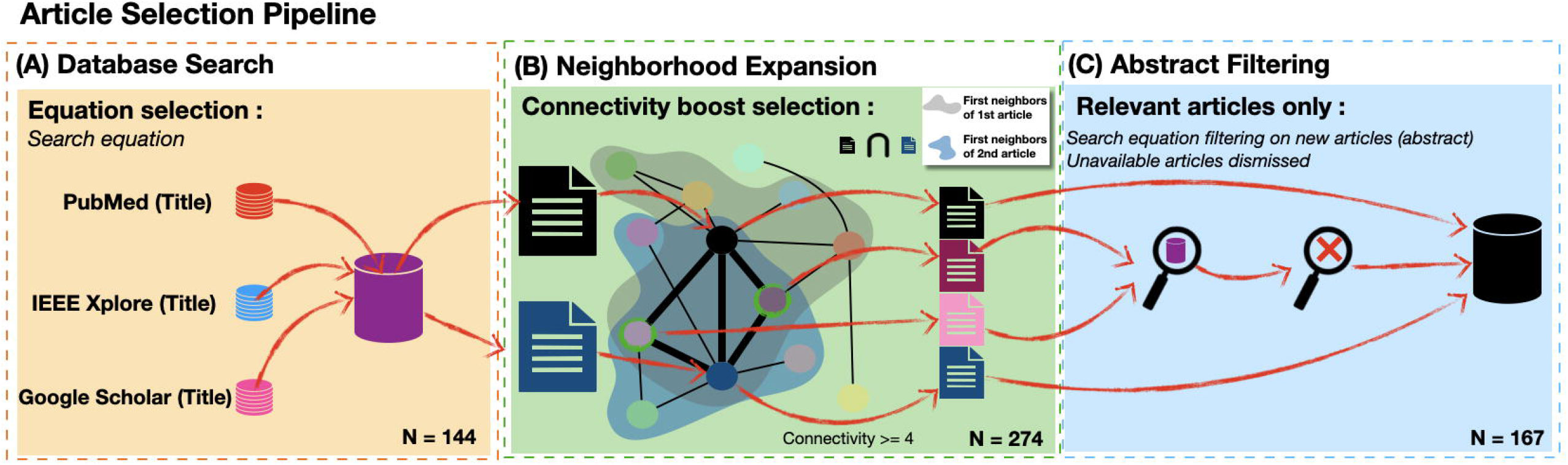
The pipeline of our methodology for choosing our articles. It consists of three steps. (A) Articles are extracted from three databases using our research equation. (B) Remaining articles are provided as anchors to Connected Papers, which generates similarity graphs for each article. Once retrieved, the graphs are integrated in a unified graph. Articles with a certain number (that we will set to 4) of links pointing towards them are added to the selection. (C) The newly added articles are filtered using the same research equation as in step (A), but searching words in keywords and abstract instead of title. Numbers correspond to the second phase of screening Overall, a total of 167 articles were used for the review. The PRISMA-style diagram synthesizing the evolution of our database is illustrated in Fig 1. Supplementary statistics and figures can be found in Supplementary Material (**Table S1**, **Table S3**, **Table S4**, **Table S5** and **Table S6**).

The Google Scholar query identified 76,300 articles. To ensure that we find as few irrelevant papers as possible, we decided to explicitly search for our keywords in the articles’ title. Although, this may seem as a drastic choice, it guarantees that we have almost only articles related to the two areas of interest. This allowed us to identify 142 relevant articles. The PubMed query applied to the title identified 56 articles, while the IEEE Xplore query identified 20 articles. By removing the duplicates, we obtained 140 unique articles after this screening step.

### Automatic enrichment with the Connected Papers tool

With such a strict search equation, it seemed appropriate to expand the set of articles identified to ensure that no critical articles were left out. To do so, we used the Connected Papers software (https://www.connectedpapers.com/), which takes an article as a query and searches a database to select the most closely related articles using a similarity measure based on co-citation and bibliography. This database is enriched with the Semantic Scholar database, which is composed of more than 240 million papers. This corresponds to step (B) of our pipeline). An example of a graph generated by Connected Papers can be seen in **Fig S1**.

We created a co-citation article directed graph for each article. 24 of them were not available on Connected Papers, or did not have enough co-citation neighbors to build a graph. This process allowed us to fetch up to 2443 new articles that were not captured by the restrictive search described in step 1 of the pipeline. For each connected-papers graph the raw list of articles was obtained. We then developed a small Python program to parse and analyze the graphs by computing various stats, including the number of times an article appeared in all graphs, along with the distribution of the number of citations (https://github.com/CorvusVaine/analyzing_connected_papers_articles.git). Finally, we computed an integrated graph, including all articles present in the Connected Papers database. The connectivity of an article in this graph varied from 1 to 34.

We decided to add to our dataset the articles with a co-citation connectivity *>* 4. We chose this threshold because it allowed us to reject as few articles as possible while not adding more articles than the original database size. For example, given a total of 144 articles, a threshold of three identifies 260 additional articles, almost tripling the dataset of articles. We therefore chose a threshold of four, which added 130 previously unseen articles. Another method tested in a first screening of articles consisted in separating them according to their subject and computing a graph for each of the different groups. This methodology can be consulted in **Supplementary Material**.

### Filtering new articles

Among the newly discovered articles, it is important to discriminate the ones that are relevant to the subject. Some of them may be important in either metagenomics or DL independently and thus have high co-citation connectivity without addressing the other topic. We thus decided to reuse our search equation as a filter for these articles, but this time by searching for keywords in the abstract and the article keywords instead of the title (See step (C) in **Fig 4**). After filtering, 23 supplementary articles are kept and added to the initial corpus for further analyses.

## Results

Metagenome-based disease prediction can be decomposed in two steps, corresponding to two scales, and therefore DL methods mostly focus at the read/sequence level and at the abundance matrix level. In **Sub section 3.1** and **Sub section 3.2**, we review sequence-based methods, respectively methods concerning functional annotation and profiling of a metagenome directly from the sequenced raw reads or generated contigs. Finally, in **Subsection 3.3**, we review the methods used for phenotype classification.

As stated before, two types of sequenced data can be seen, shotgun sequencing or 16S sequencing. The latter one produces a very smaller amount of sequences. This is why this data source is used by methods that focus on speed, such as [51] and [52], or methods that aim to create fixed pre-trained embeddings, like [53]. However, today, most methods for metagenomic analysis rely on Next-Generation Sequencing.

### Functional annotation

Next-Generation Sequencing have created a large amount of short and long reads. If classifying reads is useful for disease prediction because it allows building a “portrait” of a metagenome, identification of sequences is fundamental to understand their roles. Here, rather than discriminating every sequence, these methods are aimed at finding those that correspond to specific categories. As an example of application, we can cite the importance to discriminate viral sequences, Antibiotic Resistance Genes (ARGs) or ORFans, etc, from the rest of the metagenome.

#### Annotation using priorly known reference features

The first of these methods aim to use DL fed with known characteristic features about the type of sequences that must be annotated. These features are prior knowledge and serve to train the network to discover sequences. We can cite DeepARG ([54]) or Meta-MFDL ([55]), which classify respectively whether a given sequence is an antibiotic resistance gene or a gene fragment. These models do this by using characteristic genes and ORF features. These features can be ORF coverage, amino acid or codon frequencies, and Z-curve, and form a vector that is then fed into a deep stacking network. This network is based on stacking successive blocks of layers that take as input both the features processed by all previous layers and the raw input. In the same way, the ONN method ([56]) uses extensive information from ontologies to build an ontology-aware Neural Network for gene discovery.

#### Research from raw reads classification

The following methods aim to classify whether sequences play a specific role. However, here most of the feature extraction process is performed using the NN rather than relying on prior knowledge. These models encode sequences so that a neural network can easily process them. One of the commonly used techniques is One-Hot Encoding of a sequence (or other derived approaches, such as a mapping of *{*A,T,C,G*}* to *{*1,2,3,4*}*). They consist in representing the sequence as a matrix of 4 (one for each base) by its length, with ones corresponding to the presence of a base at a given position. These sequences are then analyzed by a neural network, which ultimately classifies them. An example is shown in **Fig 5**

**Fig 5.**
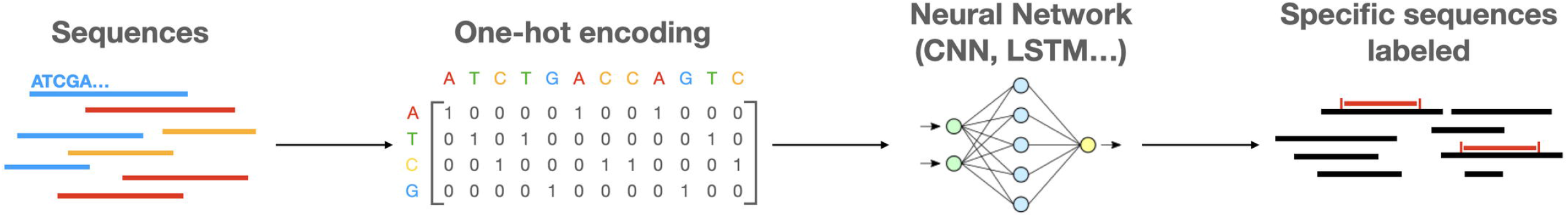
Sequence mining workflow diagram. DNA sequences are encoded, most of the time with *one-hot encoding*, which leaves a matrix of dimensions 4 by the length of the sequence. The sequence is then analyzed by a neural network, often a CNN, to be classified as a specific type of gene, for instance a viral sequence. (adapted from: [57])

This is the case of CNN-MGP ([58]), which uses a Convolutional Neural Network to extract patterns from a one-hot representation of an ORF (opened reading frame) and classify it as a gene or not. This method also allows differentiation between host sequences and those coming from the microbiome. Several methods search for plasmids and phage sequences among metagenomic sequences: tools like PlasGUN ([59]), PPR-Meta ([57]), and DeephageTP ([40]) achieve better performance than alignment-based methods in detecting phages and plasmids by one-hot encoding DNA sequences and/or proteins and analyzing them with CNNs. The latter in particular outperforms VirFinder ([60]), a virus identification method that has now been adapted to a DL architecture. In fact, DeepVirFinder ([61]) was developed using a similar approach (one-hot encoding and convolution). RNN-VirSeeker ([62]) relies on encoding sequences but considers a sequence as a temporal series and therefore analyzes it temporally using a recurrent neural network ([39]). Although trained on long reads, it performs better on short reads than previous methods because it captures the sequential nature of DNA rather than local features, changing the analysis paradigm. To date, CNNs show the best performance in this type of sequence classification problem.

Some tools, also designed to identify viral sequences, now use more than simple sequence encoding, counting on deeper features. These methods, represented by CHEER([63]) and CoCoNet([64]), rely on k-mer embedding and computed features (here, k-mer distribution and coverage), respectively. These features, which we will specify and develop later, allow them to achieve state-of-the-art or even better results in viral sequence classification. This is the reason why they are widely used.

#### NLP-based analysis

In the last few years, a new paradigm has emerged in the analysis of metagenomic sequences, very different from those previously covered. They are based on the recent breakthroughs in Natural Language Processing (NLP) using attention, Word Embeddings and Transformers, and are to applied to DNA. These methods are used to model the meaning of a text by representing various units of a sentence as mathematical vectors. DNA also has its own alphabet with nucleotides, sentences with sequences and even possibily words with k-mers. This analogy opens the way to analyzing DNA by adapting NLP methods.

Various methods use sequence embedding techniques to embed their sequences. MetaMLP ([65]), for example, embeds k-mers with a small alphabet and partial matching. This fast method allows for rapid functional profiling. DETIRE ([66]) uses methods close to the ones seen before, but by combining one-hot encoding with TF-IDF embedding of k-mers for virus detection. The structure of the data is also captured with a graph that links k-mers to their original sequences and their label (viral or not). Finally, CNN and LSTM layers aim to capture both spatial and temporal features. Virsearcher ([67]) also uses word embedding and CNN to analyse the sequence and combines the output with hit ratio of the virus.

Although these methods use word embedding techniques, new DL methods exist using the mechanism of attention.

Attention-based tools and in particuliar transformers are quite recent, but their application seems well-suited for sequence classification. VirNet ([68]) uses a deep attention model to perform viral identification and achieve SOTA accuracy. Famous Transformer models have also been adapted here: ViBE ([69]) uses a hierarchical BERT model to classify viruses at order level by pre-training it with reference virus genomes. It outperformed alignment-based methods. [70] also adapted BERT models for anti-microbial peptides identification. Finally, DLMeta ([71]) combines both CNN and Transformer to capture both local and global features from sequences. This allows to perform various metagenome identification tasks such as viral identification, but also gene prediction or protein domain prediction.

### Sequence grouping: from reads to metagenome profiling

Here, rather than identifying the type or function of a specific sequence, we focus on methods that allow to group sequences/reads into bins and subsequently profile a metagenome (see **Introduction**). Many non-DL based methods have been developed to perform such tasks and show impressive results. Many of them allow to bin contigs into genomes and thus provide a list of species representing the microbiome. We can cite MetaBAT ([72]) and MetaBAT 2 ([17]), which use probabilistic distances and tetranucleotide frequencies, as MaxBin ([73]) and MaxBin 2 ([74]) do. Finally, a method like GraphBin [75, 76] use assembly graphs and De Bruijne graphs to cluster contigs. On the other hand, the method described by [77] uses ML to compute taxonomic classification of metagenomic sequences. All of these methods provide good results when binning natural and synthetic datasets, such as CAMI datasets ([78]). But DL methods bring numerous novelties notably in terms of discovering new relevant features and embedded representations.

#### Composition-based methods

The one-hot encoding of a sequence is a limited method with respect to the goal of grouping it with others. It should be noted that various methods perform binning using autoencoders but relying on one-hot encoding ([79] and [51]) or reference database annotations only ([80]). However, these methods are now outperformed. Indeed, other features can be extracted from a sequence that better represent it. For instance, several methods work with what can be called *computed features*. This means that they process a sequence by modifying its representation with features inferred from the reads. In particular, tetranucleotide frequencies ([81]) and, more generally, k-mer frequency distributions are well known for their utility in characterizing sequences, acting like “signatures”. We will refer to these methods as “composition-based methods”. The best results are obtained using 4-mers, which corresponds to TetraNucleotide Frequency (TNF). Their distribution is computed and results in a vector representing the sequence (In the case of 4-mers, as reverse-complements are considered as one, this vector is of length 136).

#### Classification of reads

Computing an abundance matrix by grouping reads taxonomically is a difficult task as reads are often quite short (100-150bp). Two paradigms can be distinguished in order to perform this quantitative analysis. The first one is direct sequence classification and therefore relies on known classification mechanisms. Here, a read is processed individually, features are extracted and then used to classify the sequence into a given taxonomic group at a certain level. These methods are often based on pre-computed taxonomic ranks and use different architectures as direct classification (classifying directly at a given level, [82] and [77]), or by using a hierarchical classifier to distinguish, for example, first at the kingdom level, then using this result to classify at lower taxonomic level until the family level, then the species, etc… ([83]). These models are used to classify sequences one at a time, their training is performed by treating the sequence features through various layers, ending with a classification layer (for example a softmax). Due to the variety of data, there is often a possibility of rejection of the read, that is too difficult to analyze. Once the classification is done, the loss is computed and back propagated through the layers cited above.

The second approach, instead of categorizing each sequence at a specific taxonomic level, employs sequence embedding and a latent space. The model processes the features of the sequences to formulate an embedding vector. This vector is then projected into a latent space, thereby producing a novel data visualization. The latent representations of all sequences constitute a spatial distribution of the data, with each point representing an individual sequence. These points can be grouped through clustering algorithms such as k-medoids or k-means ([84] [80]). Once clustered, these sequences aggregate into groups representing their proximity in the embedding space, and therefore hopefully their real proximity. These groups and their population will form the abundance table. An example of such a pipeline is given in **Fig 6**.

**Fig 6.**
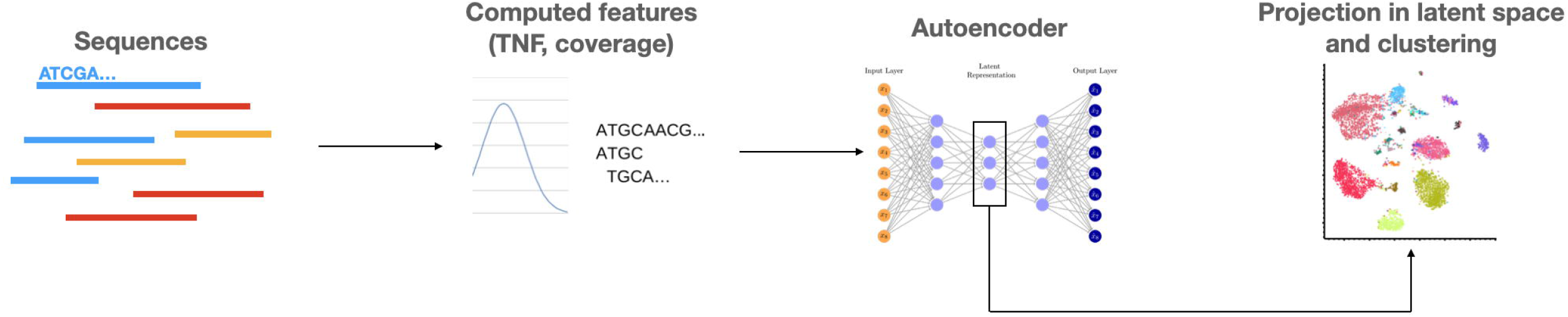
Example of an unsupervised binning method using autoencoder. Features like TNF (TetraNucleotide Frequency) or coverage are extracted from sequences and analyzed by an autoencoder, to create an embedding vector representing the sequence. This vector is then projected in a latent space, allowing visualization and clustering of sequences. Adapted from [85]

Different DL architectures can be used to embed this distribution into a vector. To extract features, methods like CNN can be used for taxonomic classification ([82], [52], [86]). Autoencoders are also useful to extract representative features from data. For instance they are used by MetaDEC ([87]), which groups reads together by creating a graph where the nodes are reads, linked if they exhibit significant overlap in their substrings. Subsequently, clusters are extracted from this graph. It then selects a subset of representative reads for each cluster of non-overlapping reads. The k-mer frequency of each subgroup is then used to build representations using autoencoders. These DL methods outperform previous methods of clustering based on dimensionality reduction, such as Principal Component Analysis (PCA), t-distributed Stochastic Neighbor Embedding (t-SNE) or Uniform Manifold Approximation and Projection for Dimension Reduction (UMAP) ([88]). They are also very useful when compared to classification methods that work with one sequence at a time because they allow to visualize the data partitioning and are therefore much more interpretable.

#### Contig binning and genome assembly

As discussed in the introduction, the primary goal of binning methods is to reconstruct the genomes of the species present in the metagenome and not necessarily to estimate an abundance table. But, by using the groups of contigs and the reads aligned to these contigs, it is possible to estimate the abundance of the species of a metagenome. In the context of contig binning, the VAMB method ([89]) was shown to outperform other metagenomic binners like MetaBAT2([17]) or MaxBin2([74]), whether for the task of classifying contigs from different types of microbiomes from simulated CAMI2 datasets or the discovery of new closely related strains. VAMB works with contigs and takes as input both the k-mer frequency and the abundance of reads mapped to the contig. These inputs are treated by a Variational Autoencoder, creating a new feature vector then mapped to a latent space. This space is then clustered using an iterative medöıd algorithm.

On the basis of the VAMB architecture, various methods have been developed for its extension or the use of other sources of information.. First, the authors of CLMB ([90]) took into account the noise, rarely considered in metagenomic analysis. To do so, they simulated different types of noise, augmenting contig data with noised sequences. The model was trained with the double objective to minimize the reconstruction error between noised versions of a same contig while spotting the differences between different contigs. This approach was based on the principles of *contrastive learning* ([91]). The network learned to handle noise by itself and pulled together in the latent space different versions of a same contig while pushing away other ones. The latent space clustering remained similar to that of VAMB. Compatible with other binners, CLMB was more refined and outperformed them (MaxBin2, VAMB and MetaBAT2) on the same CAMI2 datasets. AAMB ([92]), an extension of VAMB, is also based on its architecture and compatible with it. Instead of Variational autoencoders, it relies on Adversarial Autoencoders. The strategy is to use the same input as VAMB and to encode it in two latent spaces: one is continuous and the other categorical. These two spaces are clustered, and a discriminator for each space makes sure the encoding stay close to its prior distribution.

Also based on Variational Autoencoders, CCVAE([93]), aims to get beyond local sequence features by taking into account for binning not only the contig itself, but also the reads composing it. To do this, they use the assembly graph, a graph where nodes are the contigs and edges the k-mers connecting contigs, with a weight equal to the number of time this k-mer occurs in the data. This graph constrains the VAE to represent nodes with edges between them with more similar features. Taking into account this graph allows this method to outperform VAMB, and paves the way to graph embedding methods in metagenomic binning.

Finally, another method outperforming VAMB is SemiBin ([85]), which follows the concept of Semi-supervised learning, by adding information from reference databases while still being able to discover new bins outside of reference datasets. SemiBin relies on the notion of constraints by creating must-link and cannot-link constraints between contigs. The must-link constraints are created by breaking contigs up, while the cannot-link constraints use reference contig annotations. These constraints are combined with the same inputs as VAMB (abundance by mapping and k-mer frequencies). Deep Siamese networks embed these features in a distance between two contigs, generating a sparse graph clustered with k-means algorithm. SemiBin outperforms existing binners, in particular VAMB and SolidBin ([94]), on both real and simulated datasets. More specifically, it recovers with great completeness a high number of complex bins. It is precise enough to differentiate *B. vulgatus* from human and dog gut microbiomes.

Noteworthy, these binning methods work with contigs rather than raw reads. Contigs must first be generated with an independent software ([95]). In particular, semiBin demonstrates the importance of background knowledge, showing the importance of continuous database progression in the binning task. To date, *sequence-composition and feature abundance* methods provide the most convincing results for this kind of tasks, but other tools use different approaches based on promising new architectures.

#### Methods inspired by Natural Language Processing

As NLP was used for functional annotation, it is also more and more used to classify reads and perform binning, or even analyse a metagenome. The metaphor is to think of each genome in the metagenome as a text, whose sequences would be sentences made up of k-mers or other features, which would be words.

DeepMicrobes ([96]) highlighted the importance of k-mer embedding. It compared this method to one-hot encoding and introduced attention in metagenomic analysis by presenting an architecture using LSTM and Self-attention based models. The results show that embeddings significantly improve performance when compared to one-hot encoding. This work has paved the way for the use of self-attention for the representation of DNA sequences, but also to the importance of their sequential nature with the use of LSTM networks. This approach performs well with long reads, but faces difficulties with shorter reads.

Given the analogy between NLP and DNA analyses, it is not surprising to see adaptations of word embedding algorithms to DNA sequence data. The word2vec method ([97]) has been adapted to generate k-mer and sequence embeddings by both NLP-MeTaxa ([98]) and FastDNA.([99]). FastDNA was reused within the Metagenome2Vec method ([100]) to combine word embeddings with taxonomy and create a metagenome embedding. Metagenome2Ve relies on such metagenome embedding to perform disease prediction. In the context of Metagenome2Vec, the term *End-to-end* implies that the method encompasses the full spectrum of processes needed to convert raw metagenomic data into valuable vector representations: inputting raw reads, quality control, feature extraction, all the way to dimensionality reduction, eliminating the need for manual preprocessing or feature engineering. Meta1D-CNN tries to enhance the precision in sequence classification with NLP methods by introducing 1D-CNN. They train a word2vec algorithm with different k-mer lengths from 1 to 8 (8 giving the best results). The embedding of a sequence is obtained by calculating the mean of all k-mer embeddings. Two layers of convolution are then used to extract features that will ultimately be used to classify sequences. On its dataset, Meta1D-CNN achieves 83.89% F1 Score at the genus level and 65.65% at the species level, while NLP-MeTaxa achieves respectively 83.89% and 60.95%.

While these methods are proof of concepts, they have not outperformed alignment-based methods outlined earlier. These Deep Learning approaches have allowed to gain insights on the limitations or difficulties with the NLP approach. First, the amount of noise in the data must be taken into account, particularly here, where sequence representation is the heart of the work. Secondly, the comparison of genomic reads to text does not fully hold up due to the intrinsic differences between k-mers and words. K-mers do not only overlap but also form a finite, known, and extremely dense vocabulary, particularly for a smaller value of *k*. Furthermore, a larger *k* value results in more accurate classification as the number of distinguishing k-mers becomes increasingly prevalent. A significant limitation of this approach is that each increment of 1 in the value of ‘k’ quadruples the size of the vocabulary. Consequently, this exponential increase leads to substantially higher computational demands.

Several ideas have been explored to solve the issue of increasing computation time with longer k-mers. One idea is to enlarge the vocabulary by taking longer k-mers, but regrouping some of them based on proximity criteria. META^2^ [101] regroups k-mers using Hash Embedding or Local Sensitivity Hashing. Reads falling in the same bucket share the same embedding. On the other hand, fastDNA has been enhanced with BRUME [102]. The idea is that k-mers that are always present or absent together in the same reads should be considered as having the same importance in sequence embedding. Therefore, they can be grouped together, using methods such as de Bruijn graphs. The *de Bruijn graph* of a set of sequences is a graph where each vertex is a k-mer and edges between vertices represent k-mers that are adjacent in the set of sequences. The graph can then be compacted along non-branching paths, creating contigs to which each k-mer belongs, and are then assigned to the same embedding. The drawback is that some k-mers present in new sequences to be analyzed may not have been seen by the network during training and have no embedding, and this becomes more likely as k grows. This methodology facilitates analyses with ‘k’ values exceeding 30, a value made possible as the quantity of de Bruijn contigs tends to plateau. The increase in ‘k’ value enhances the effectiveness of this method, thereby leading to better results.

These ideas open the way to new methods using in metagenomic binning recent NLP methods such as BERT ([103]) and its successors. Several studies have attempted to adapt the BERT method to metagenomics, but because these models are computationally expensive, they have not gone as far as they could to produce usable results.. Bi-Meta ([104]) adapts various NLP techniques (Latent Dirichlet Analysis (LDA) or Latent Semantic Analysis (LSA)) or models (Word2Vec and a very small version of BERT), while BERTax ([105]) also tries to train a small BERT model to perform taxonomic classification of sequences. It reproduces the masking process but uses non-overlapping words instead of k-mers. The results of these models show that although BERT is a very powerful model, especially in detecting sequences that are not closely related, it is still limited by both its computational cost and the large diversity of microbiomes. This diversity is not yet well represented by the available data that these models would need for pre-training to achieve better performance.

A recap of methods dealing with phenotype prediction can be found in Table 1, and some performance comparisons can be found in Supplementary Material on taxonomic classification: Tables S1, S2, S3, S4 and S5.

**Table 1.**
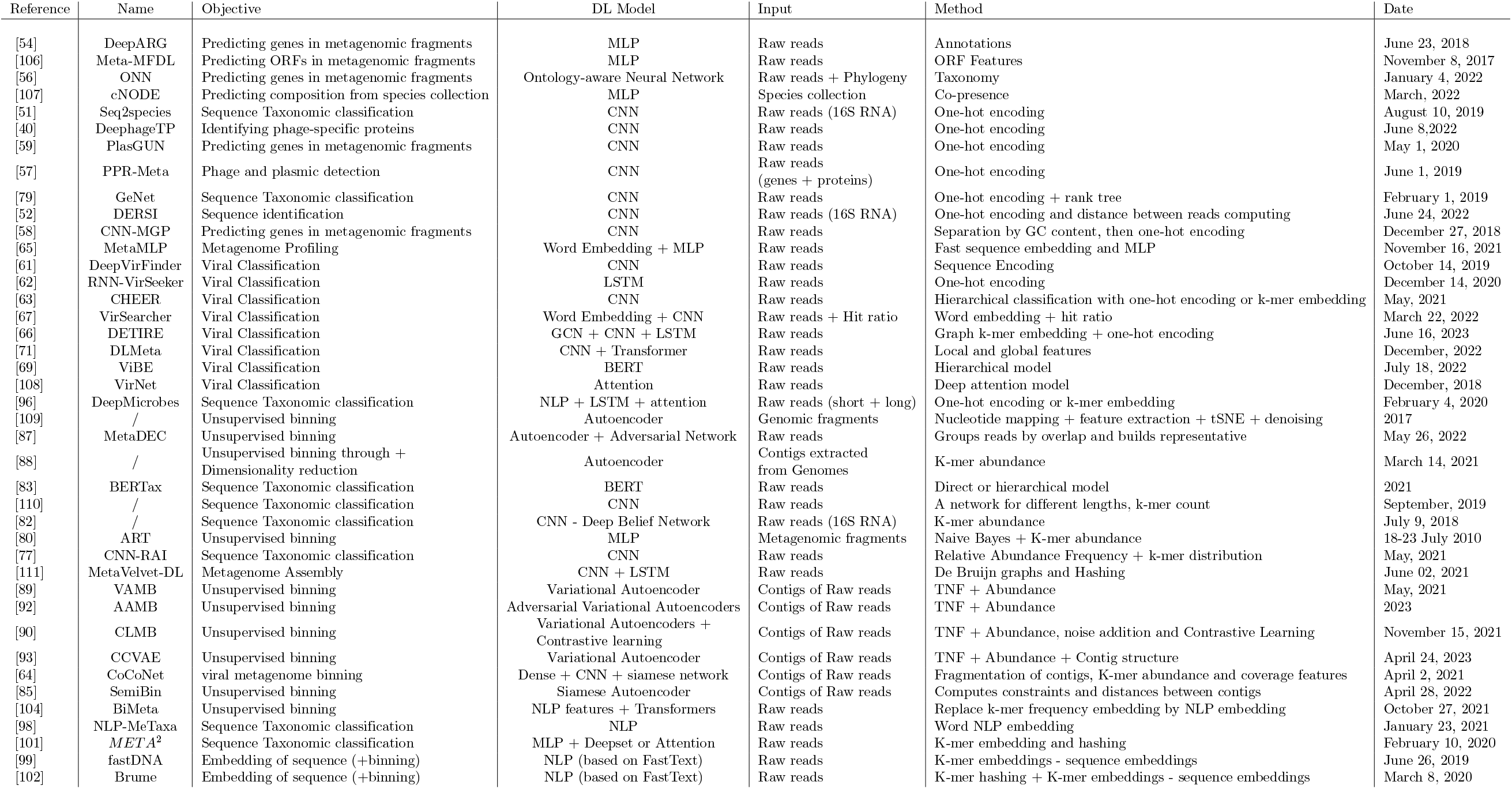
Table listing different DL-based methods as well as their performance in taxonomic classification. Please note that the results are found on each model’s own dataset, not on a centralized dataset, using different metrics and data of various complexity, and can therefore not easily be compared.

### Phenotype classification

The binning process itself is quite important, but in the case of disease prediction, it can also serve the purpose of microbiome characterization and quantification, which is useful for extracting metagenomic information and using it for diagnostic purposes. Machine Learning has already shown its effectiveness in diagnosing disease from metagenomic data. These approaches include MetAML ([112]), Predomics ([48]),SIAMCAT ([113]), etc. Diseases are not the only characteristic which can be inferred from metagenomic data: [114] for example does not perform disease detection, but tries to predict an individual’s age from their microbiome using DNN. This demonstrates the richness of applications of metagenomic data. Most often, what is used to classify phenotypes are abundance tables of different taxa obtained after binning. They are usually tables where the rows represent the samples examined and the columns represent the taxonomic abundances.

Metagenomic abundance data are sparse and the number of features greatly exceeds the number of samples([115]), making it challenging to train models that do not overfit. There are several solutions to this problem including data augmentation.

#### Data augmentation

Despite lowering costs in sequencing data over the past decade, data accessibility still remains an issue, particularly with regard to the availability of metadata (especially clinical patient information). Besides real data, it is also possible to simulate metagenomic data using simulators such as CAMISIM ([78]). Other methods deal with the problem of unbalanced classes by oversampling ([116]). The method developed by [117] handles this by resampling the poorly represented class until they all have as many samples as the most represented and reweighting each class, then training the classifier in a one vs all manner for each of them.

For the specific case of metagenomics, [118] have proposed a DL based approach to address this issue by generating new samples using Conditional Generative Adversarial Networks (CGAN). The idea behind a GAN is to use two competing networks: one to generate data coherent with the input dataset, and the other to try to detect whether that dataset is real or generated. The two models are trained in an adversarial way. CGANs offer the possibility to parameterize this generation: the network can then decide to generate for example healthy or disease-related data. But the issue with GAN is that finding an optimal model is often challenging, and therefore there is a risk of generating unrealistic data. Furthermore, their training requires a large amount of data. Although the proof of concept is promising, it is still a problem to get sufficient quality data to train GANs and subsequently classification models.

Another method that relies on DL is explored by [106] who use Variational Autoencoders to generate new data. It works by reconstructing modified versions of their original metagenomics data. Variational Autoencoders use the probability distribution of the input data to generate new modified samples. [119] also used this approach combined with a chained normalization method and feature extension. Methods such as MetaNN [120] show that it is possible to achieve better classification results compared with classic ML methods using simple NN and data augmentation.

The value of data augmentation lies not only in the fact that it lowers overfitting, but also in the fact that it addresses problems related to unbalanced data sets. [117] is confronted with this problem in particular when trying to build a multi-class classifier of 19 diseases from 5 different body sites. Using class weighting and resampling, it achieves results that are inferior to the state of the art, due to the number of classes, but which can rival conventional methods when it comes to evaluating not just the Top 1 predicted diseases, but the Top 3 or Top 5, despite a highly diverse dataset. On the other side, MegaD [121] is a simple Neural Network specifically trained for large datasets that achieves competitive results while running relatively fast on large datasets.

#### Abundance-based approaches

As mentioned above, microbiome abundance data is not well suited for direct Neural Network analysis. Therefore, how the data is used and fed to DL tools is crucial to extract meaningful representations of the input. This can be done with Deep Learning representations.

#### Learning new representations

To deal with the issue of high number of features in metagenomic data, many methods use dimensionality reduction techniques. These methods consist in representing very sparse data in a smaller dimension, reducing the imbalance observed before. It is possible to use different feature selection methods as well as DL-based data transformation methods.

#### Mathematical Transformations and Feature Selection

Mathematical transformations of data are a good way to generate novel representative features of the input, here the abundance tables. Following this idea, [106, 119] use different normalization methods combined with autoencoders to extract exploitable features.

These mathematical transformations create new representations from the data that are easier to use by DL. Many feature selection methods have been used to reduce data sparsity. [122] uses Ridge Regression algorithm on Gene Family Abundance to create lower dimension data that can then be analyzed with a CNN.

While most data preprocessing methods use normalization or distribution algorithms on input tables, [123] bypasses the DL training step by directly using statistical binning methods such as Equal Frequency Binning or Linear Discriminant Analysis, and K-mean Clustering after that. This work directly bins metagenomes and associates them with the correct disease, achieving good prediction accuracy.

#### Reducing dimension through autoencoders

Since the extraction of relevant features is a specificity of Deep Learning, different types of NN have also been used to obtain better representations and embeddings. The main issue encountered with feature selection is the loss of potentially important information. It is therefore of great importance to find efficient dimensionality reduction methods. As stated by [124], autoencoders are an interesting hypothesis offered by DL for relevant task-adapted dimensionality reduction. Indeed, autoencoders are known for their ability to extract relevant data representations with dimensionality reduction using the canonical encoder-decoder architecture. Such architecture is well suited to deal with the problem of sparse matrices and low sample number. Moreover, training of the autoencoder causes the data reduction method to be adapted to the specific structure of the data. The newly generated features can then be used for classification by classical Machine Learning methods.

However, s the best type of autoencoder to use remains still an open research area. For example, DeepMicro [125] chooses to train different types of autoencoders to find the one that extract the most significant information for disease prediction from metagenomic data. Sparse Autoencoders (SAE), Denoising Autoencoders (DAE), Convolutional Autoencoders (CAE), and Variational Autoencoders (VAE) were all tested and gave good results, none of them outperforming the others, as the best result depended on the datasets which contained data corresponding to 6 different diseases. The conclusion drawn by the article was that the best type of autoencoder depends strongly on the structure of the data (CAE when working on Abundance profile for example, while none outperformed the others on Marker profile) and that there is no absolute good answer. However, due to the black box nature of the models, the difference mostly lies in the performance in predicting several diseases (DAE for Obesity and Colorectal, and CAE for C-T2D and Cirrhosis) and does not offer much insight on the reasons why some autoencoders perform better than others.

Ensdeepdp takes into account the fact that different types of autoencoders seem to extract various useful types of features by using Ensemble Learning to get the best possible representation ([126]). Based on the idea that autoencoders should have difficulty reconstructing metagenomes from patients suffering from a disease as they can be quite different from the healthy ones, it focuses on reconstructing them using autoencoders. The distance vector between the original metagenome in input and the reconstructed one in output acts as a disease score. This experiment is repeated with many Autoencoders, VAE and CAE, with different architectures and parameters. A parameter of k is introduced and the k best models are then selected. When analyzing a new metagenome, a matrix composed of the input data and the k best models’ representations of this input data are computed, thus enriching the original feature space. The new representation of the metagenome is then composed of the original abundance vector and the newly generated features of the best models. Essentially, the approach introduced here involved training multiple deep learning models on the metagenomic data, and then combined the predictions of these models to make a final decision about the presence or absence of a disease. The goal of this approach is to improve the accuracy of disease prediction compared to using a single deep learning model.

#### Pretrained matrices of metagenome embedding

Some methods propose pretrained tools that rely on NLP mechanisms to generate embedding matrices that can then be reused with new data. Once the matrix of embeddings is created, the new data is simply multiplied by the embedding matrix to produce a new table of embedded data. GMEmbeddings ([53]) provides embeddings based on GloVe ([127]), a Natural Language Processing algorithm, by aligning requested samples to known ASV using BLAST. ([128]) uses the same GloVe algorithm to generate an embedding of a user-uploaded abundance matrix. The newly created data embeddings can subsequently be categorized using traditional Machine Learning algorithms, such as Random Forest.

### Sequence-based approaches

#### Sequence embeddings

While most phenotype prediction methods rely on taxonomy and abundance, some use other sequence-based features. They learn embeddings of relevant sequences to classify directly with them, or to enrich abundance and composition data. These approaches have the great advantage of being “end-to-end”, they can avoid the computational cost of binning methods, alignment-free or not, or use binning as an auxiliary source of information.

We have already emphasized the efficiency of k-mer distribution analysis for binning. K-mer distribution also proves useful for prediction. MicroPheno ([129]) is based on the k-mer distribution of shallow sub-samples of 16S RNA sequences. A bootstrapping framework selects relevant sequences before computing k-mer representations, allowing classification and visualization of important sequences. Aggregation of these representations allows phenotype prediction. However, the problem with such aggregation is the loss of information over microbial interactions. K-mer distribution based embedding is then compared to another method using learnt embeddings by ([130]). Here, the embeddings are discovered using the NeuroSEED framework ([131]), which uses an autoencoder to compute the distance between sequences. This allows to represent each sequence in a latent space when compared to each other.

However, instead of the distance between sequences, another analogy can be considered for metagenomic data. This analogy is that of natural language and its connection to the language of DNA. K-mers are compared to words, sequences to sentences, and metagenomes to books in order to adapt word integration architectures to the task. We have already discussed this analogy concerning the task of sequence binning. The idea here is to embed reads and use these embeddings for disease prediction. For example, IDMIL ([132]) uses bag-of-words TF-IDF algorithms to obtain an embedding for each k-mer. It aggregates these k-mer embeddings to get read embeddings. Using the same idea, Metagenome2Vec ([100]) avoids the solution of simply aggregating data, which would lead to losing precision, by using fastDNA ([99]). Using FastDNA on metagenomic data, it performs both read embedding and read binning, taking into account the link between words and sentences, here with k-mers and sequences.

#### Multiple Instance Learning with sequence embeddings in Prediction

Metagenome2Vec ([100]), IDMIL ([132]) and the method described in [130]) use a particular DL paradigm called *Multiple Instance Learning*. Multiple Instance Learning (MIL) is a supervised learning paradigm that consists of learning from labeled sets of instances, known as ‘bags’, instead of learning from individually labeled instances. Each bag is associated with a single label, and contains multiple instances. ([133]). The fundamental assumption in MIL is that a bag is labeled positive if at least one instance in the bag is labeled positive. If they are all negative, then the bag is labeled negative. Some methods have used this paradigm to perform phenotype classification from raw sequences instead of abundance tables. When using abundance, the information carried by a sequence is reduced to the species it belongs to. With MIL, it is possible to represent a metagenome as a bag of sequence embeddings, thus keeping the information of the sequence. However, each metagenome contains millions of sequences, which represent a gigantic computational cost. Therefore, most of the time, not all sequences are treated, but rather groups or representatives of sequences.

In the method from [130], sequences are represented through NeuroSEED. As they are obtained from 16S data, there are notably fewer sequences. They can therefore use the whole set of sequences. The problem is considered as a set classification, using all vectors and not their aggregation. To solve such a problem, they use MIL architectures like DeepSets ([134]) and Set Transformer ([135]). IDMIL and Metagenome2Vec, on the other hand, use shotgun metagenomics data, composed of millions of sequences. The computational cost of studying millions of sequence embeddings by sample makes this idea unreasonable. But this computational cost can be drastically reduced if instances are not sequences themselves, but groups of sequences. An example of their pipeline can be seen in **Fig 7** This is the idea followed here, with IDMIL ([132]) where sequences are clustered by a k-means algorithm and a representative of each cluster is used, creating “instances”. These instances are then ordered following their distance to a “center”, computed by using the center of the different centers of clusters. This order creates a matrix of representatives’ embeddings, that is then analyzed by a CNN. Attention mechanism is also performed on this data. It allows to differentiate and learn about the predictive interest of a given instance in the bag for metagenomic classification: which sequences are important for disease detection and which are not. However, attention being performed before the CNN, it is quite difficult to assert that it represents the true importance of each cluster. With Metagenome2Vec ([100]), read embeddings are clustered by species through binning with fastDNA ([99]) to obtain an embedding of each taxon. These taxa embeddings are the instances that form the core of the Multiple Instance Learning method used here. The metagenome is then a bag of taxa embeddings that can be analysed with MIL architectures like DeepSets and MIL-VAE. This approach is promising and end-to-end, although it still requires a binning phase. However, the way in which embeddings are exploited remains to be improved.

**Fig 7.**
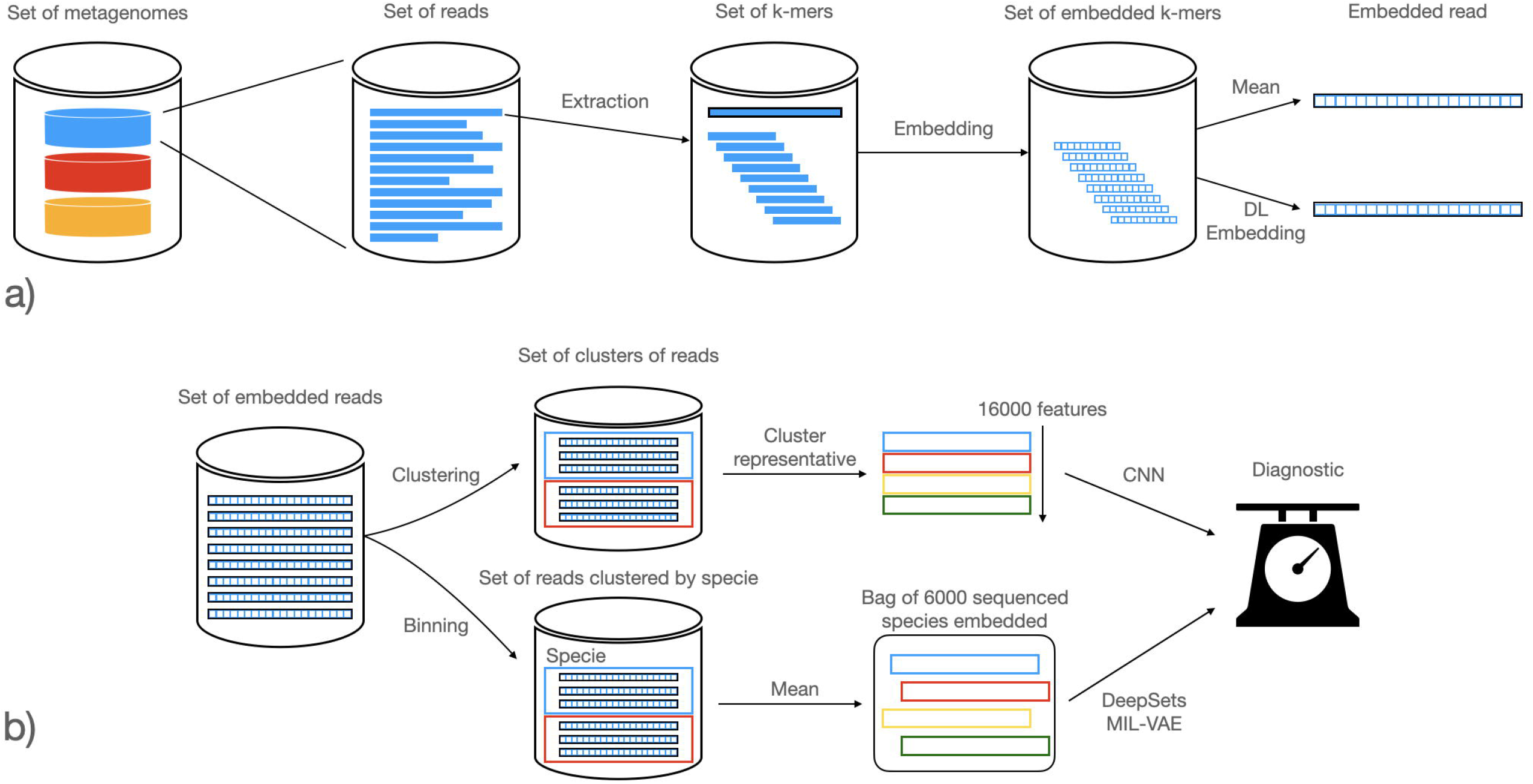
Classification with sequence embedding MIL pipelines. This pipeline is shared by both Metagenome2Vec ([100]) and IDMIL ([132]). The arrows above correspond to IDMIL, the lower ones to Metagenome2Vec. Step a) presents how sequences are embedded: their k-mers are extracted and embedded using NLP methods. These embedded k-mers are then used to obtain the embedding of a read, whether through their mean or by learning the relationship between k-mer embeddings and read embeddings through DL. Step b) presents how these embedded reads are grouped together. IDMIL uses unsupervised clustering with k-means, while Metagenome2Vec groups reads by genomes. Both obtain groups of read embeddings, that must then be embedded themselves. Here, IDMIL chooses a read representative for each group, while Metagenome2Vec chooses the mean. These group embeddings represent the metagenome differently: the first method orders them in a matrix and uses a CNN for prediction while Metagenome2Vec treats them like a bag of instances and uses MIL methods such as DeepSets ([134]) to analyze them

This paradigm, while still relatively underrepresented in contemporary literature, presents a compelling approach due to its ability to operate at a granular sequence level. This contrasts with the utilization of abundance tables, which are commonly associated with several drawbacks such as sparsity, complexities in construction, information loss, and dependency on catalogues. As such, the adoption of this paradigm could potentially address these challenges and enhance the precision and efficiency of machine learning applications in this domain.

#### Integration of other types of data

Acknowledging that raw metagenomic data is not always well suited for DL, other types of data than abundance tables can be fed to give coherence to metagenomes. They are diverse and can come from the data itself or from external knowledge.

#### Taxonomy-aware learning

Abundance tables, while providing measures at species level, do not provide information on their relative evolutionary distance. Species with close genomic sequence share similar functions and are potentially adapted to the same environment. Such information can be represented as a taxonomy tree and integrated with abundance information directly when training NN for classifications tasks. Several approaches have been tested to integrate taxonomy information: MDeep ([136]) groups OTU in its vector by using a measure of correlation structure based on distance between OTUs in the tree, hoping to make phylogenetically correlated taxa close to each other. Then, authors designed a CNN with three layers that are supposed to mimic the different levels of phylogeny and their interactions, with smaller numbers of neurons each time, supposedly corresponding to Genus, Family and Order, before using Dense Layers. TaxoNN ([137]) uses a comparable yet different technique: it groups each abundance unit according to their phylum and trains a Convolutional Neural Network for each phylum, learning the features specific to that phylum. Feature vectors from each network are then concatenated and used for final classification. The problem is then deported from species level to phylum, and phylum analyzed separately before the dense layers.

Ph-CNN ([138]) takes this idea further by using the distance measures in the taxonomic tree to take into account the proximity between taxa. A custom layer is designed to perform convolution on the k-nearest neighbors’ abundances. This method is highly dependent on the chosen distance. The drawback is that although it takes into account neighboring taxa, it focuses on local patterns and does not process the structure of the data globally.

PopPhy-CNN ([139]) proposes a tool that embeds the taxonomic tree in a matrix, allowing all the topological information to be processed. **Fig 8** shows the embedding algorithm chosen by PopPhy-CNN. This embedding is designed to avoid sparse matrices. The drawback of this representation is the structure of the matrix itself: embedding a tree in a matrix can result in very sparse matrices. To avoid that, this method places all nodes at the leftmost non-null spot in the matrix. A consequence is that, with a more complex tree and as nodes are placed to the leftmost spot, some nodes may not be found directly above their parents, thus blurring the links that the tree is supposed to represent. For example, in **Fig 8**, node labeled 5, found in coordinates (5,4), is directly under the node labeled 8 (4,4), when it is not its descendant. To consider more of the tree structure, TopoPhyCNN ([140]) embeds it in a matrix, but adds topological information like number of child nodes, height of layers and node distance in the tree.

**Fig 8.**
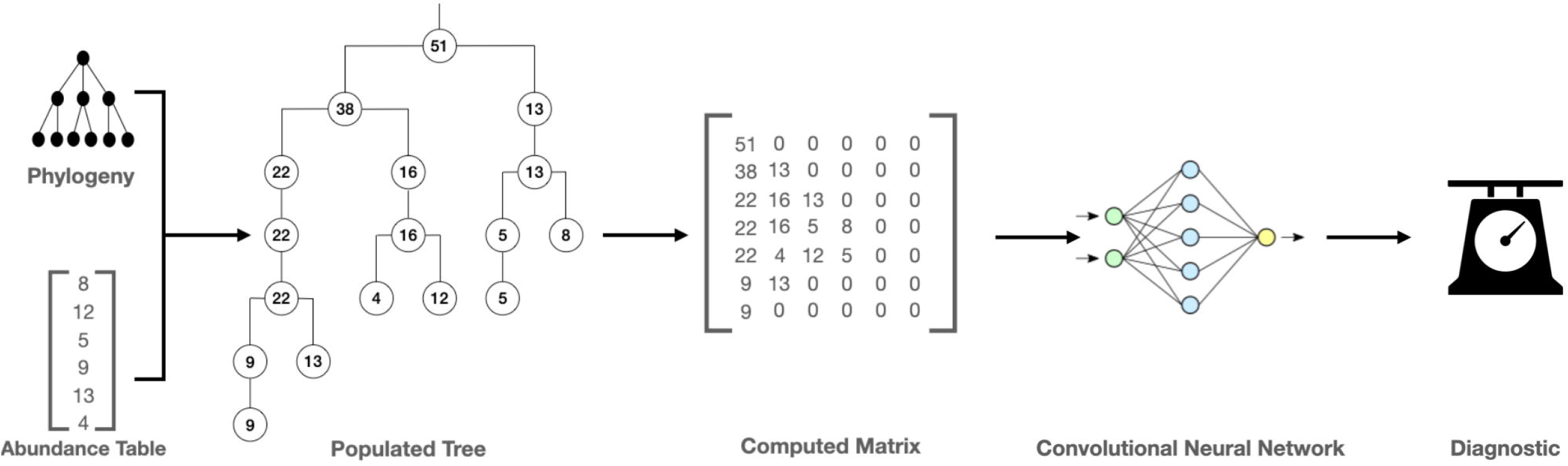
Taxonomy-aware metagenome classification method, as performed with PopPhy-CNN ([139]). Phylogeny between taxa is used to create a tree, and abundance to populate it. This tree is then embedded as a matrix used as input for a Convolutional Neural Network that will ultimately classify the metagenome. Modified from Source: [139]

**Fig 9.**
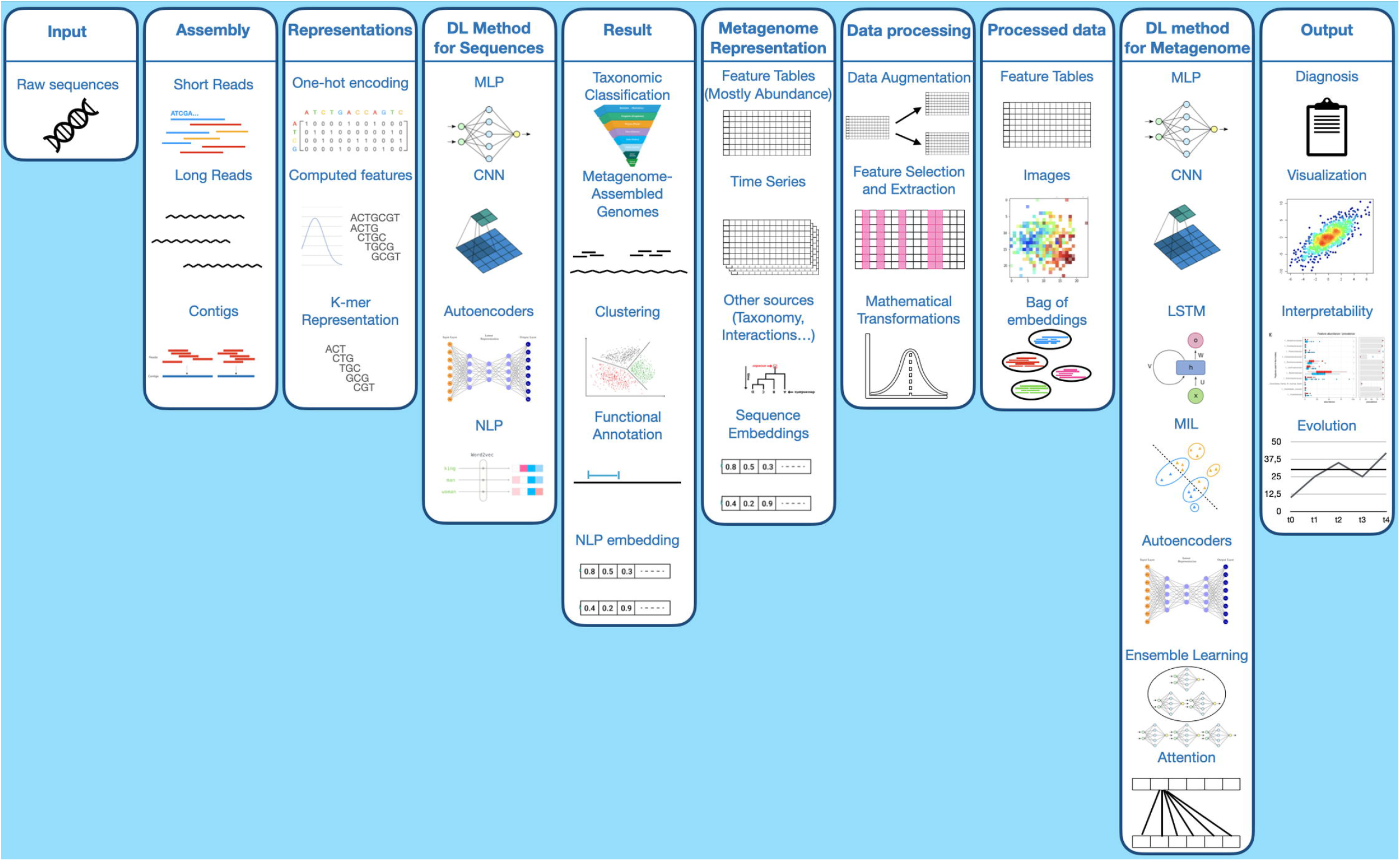
Overview of different steps and methods in disease prediction from metagenomic data. These steps represent the entire pipeline from raw reads to disease prediction. It is to be noted that not all steps are required and some methods described in a step are not always compatible with every method from the next step. This figure aims to represent the diversity of method in each step, not necessarily every entire pipelines possible. Moreover, as said before, most methods only perform one half of the steps: the first half from reads or contigs (steps **Input** or **Assembly**) to their classification (steps **Result** or **Metagenome Representation**) and the second half for disease prediction (step **Metagenome Representation** to **Output**). **Input** represents the raw sequences acquired through sequencing. **Assembly** can either be the long or short reads acquired before, or the contigs assembled from these reads. **Representations** are the way these features will be fed to the DL model (encoding, features). **DL Method for Sequences** show the different types of networks used to extract features. **Result** are the output of these networks: classification, clustering, embedding, etc, which can then be used for **Metagenome Representation**, along with other sources. These representations are then filtered of transformed through **Data processing**, resulting in **Processed data**(images, tables, clusters). **DL method for Metagenome** are then used to treat these features and produce an **Output**: diagnosis, data visualization, phenotype evolution.

These tree and graph structures present a very complex, big and potentially sparse structure. This is a serious limitation that is acknowledged by the authors, who encourage the exploration of other embedding methods. To give coherence to abundance data, some authors have tried to take spatial embedding to the level of the image: abundance data is converted and represented by an image. The Met2Img method ([36]) used this paradigm to outperform previous state-of-the-art tools. The abundance vector is represented as a 2D image, colored by a taxonomy-aware fill-up method. The generated images are then analyzed by a CNN to retrieve more structural metagenomic information. Furthermore, this method can be combined with the use of other omics or patient data.

1. [141] offers direct comparison between tree-embedding methods and new image representations to show the advantages of the latter. By taking the most represented genera, they create different types of image representations with each genus represented by a shade of grey linked to its abundance. These images can then be analyzed with a ResNet-50, a DL image analysis technique. A great advantage of this method is its interpretability, because genera that were useful for prediction of disease (here Type 2 Diabetes) can be easily traced. However, this method works at the genus level, at best, and by taking into account only the most represented genera in the data, therefore potentially omitting information coming from less represented bacteria.

Following the method of Met2Img, the more recent MEGMA method ([142]) uses Manifold Embedding to create a data embedding based on co-abundance patterns between microbes. 5 Manifold Embedding methods were tested, as well as Random-guided Uniform Embedding: MDS, LLE, ISOMAP, t-SNE, and UMAP. On the other hand, microbes are grouped based on their phylogeny. This grouping will determine the color used in the image for each group. In summary, the localisation on the image is based on the embedding, while the color is based on phylogeny, the opposite of Met2Img. This new method outperforms Met2Img and is as well very interpretable, for parts of the images important for prediction can be found and linked to the microbes they represent.

Finally, another aspect that can be taken into account when taxonomy is studied is the fact that a great part of it is unknown, whether it is because abundance is obtained by unsupervised binning or because reads come from unknown species. MetaDR ([143]) takes into account both known and unknown features as well as the topology of the taxonomy tree obtained by converting it to an image, allowing MetaDR to compete with the best state-of-the-art methods, while showing good computational speed and ranking among the best taxonomy-based methods.

#### Microbial interactions

While taxonomy offers valuable insights into the relationships between microbes, it only captures a fraction of the complex interactions within the microbiome. Microbes interact and function in myriad ways within this environment, and their taxonomic connections alone are insufficient to fully comprehend the intricate dynamics of this ecosystem. Therefore, a more holistic approach that goes beyond taxonomy is necessary to unravel the comprehensive functioning of the microbiome. ([144]) attempts to tackle this issue by using the abundance of each species to compute various sparse graphs of interactions between species using co-abundance patterns. The graphs are then fed into a Graph Embedding Network designed with a specific layer for graph embedding. Despite the interesting questions raised by these methods, finding other ways to analyze interactions between microorganisms remain under-explored in the field of DL and an issue still to be addressed.

#### Functional and genetic information

Some authors have chosen to use the functions of genes or specific communities contained in a metagenome. However, metagenomic diversity remaining largely unexplored, using reference databases might be challenging or incomplete. Still, some tools try to extract relevant information from these databases. Most of these tools rely on classical Machine Learning and not Deep Learning. [145] uses functional profiles extracted from orthologous genes given a reference database to add these features to abundance, while DeepMicro ([125]) uses strain-level marker profiles to contextualize and deepen abundance data by the presence or not of a certain strain. As for abundance data, strain-level markers is very sparse information, leading to the same difficulties. However, methods like Principal Component Analysis have shown satisfying results when applied on this data, leading to a slight improvement in prediction. The other way around, ML methods like [48] and [146] aim to extract top decisive features or markers for disease prediction to understand key roles played by these features in the apparition of a disease.

#### Combining different sources

Using Deep Learning to try and conciliate many ways of integrating information, MDL4Microbiome ([147]) opens the way to adding different types of data for prediction by designing a model made of various parallel simple feed-forward neural networks. Each network takes a different source of data as input and performs phenotype classification. By concatenating the last features used before classification of each network, MDL4Microbiome can obtain a vector representing each source. This model seems to outperform classical ML methods in disease classification, and shows that combining features together improves results than using each feature type separately. Here, the experiment is performed with three sources of data: species abundance, metabolic function abundance, and genome-level coverage abundance, but any feature can be used following this simple model, even though its use might not be optimal.

#### From cross-sectional to longitudinal metagenomics data

The human microbiome is highly dynamic and can change drastically in a short time, be it due to diseases, diet or medical interventions. All the methods described above work with single-point data. However, it is possible to study the evolution of a microbiome over time or the influence of specific events on its composition with longitudinal data, i.e. at different time steps from the same patient. [148] for instance analyzed such data before and after dietary changes to understand their impact in the microbiome composition. With the same idea, [149] studied the transition from adenoma to cancer, although their method does not use DL. GraphKKE ([150]) on the other hand, used a DL based approach and proposed to embed a microbiome with time-evolving graphs. Nevertheless, these methods are not strictly speaking temporal studies. The data is not seen as temporal series, and therefore the analyses are independent single-point analyses, and not an analysis of the evolution of the microbiome through time. The temporal study is more seen as giving coherence between different time steps and studying the longitudinal metagenomic data as a whole, rather than different time steps without linking them together.

There are other methods based on DL used to analyze real time series data. Instead of a single point abundance vector, they consider a time series of vectors, which means a matrix containing a vector for each time step. It can be done through the use of Recurrent Neural Networks (RNN) and in particular Long Short-Term Memory (LSTM) models. These networks capture the temporal evolution of data through different time steps. [151] used this method to predict the occurrence of allergies in children aged 0 to 3 years old, while [41] used them to predict the evolution of ulcerative colitis (UC) and [143] classified various diseases like type 2 diabetes (T2D), liver cirrhosis (LC) or colorectal cancer (CRC). All these methods used phylogenetic information of different time steps treated as a time serie by a LSTM. This has proven more effective than SVM, KNN or LR Machine Learning methods. To try and give more coherence to both each time step and their global dynamics, an approach combining CNN and LSTM was developed with phyLoSTM ([152]). Here, each time step is processed following the same method as with TaxoNN ([137]), ie by ordering OTU by phylum and using a CNN adapted for each phylum. Once the feature vector for each phylum is extracted, they are concatenated in a feature vector representing the time step. All these vectors will then form the new time series to be analyzed by the LSTM. Therefore, phylogenetic information is extracted by the CNNs, while temporal features are extracted by the LSTM.

This CNN-LSTM structure has also been used in [153], but enhanced with self-distillation ([154]). Knowledge-Distillation ([155]) is a recent and impressive neural network training technique. It consists in transferring knowledge from a large and heavy model to a lighter one by training it to mimic its output. This technique saves a lot of computation time, despite a degradation in accuracy. Self-distillation consists in applying such a process to a network by itself. It is done by plugging shallow classifiers at the output of hidden layers in the network. These classifiers allow to compare the features outputted by hidden layers to the global output of the model, and therefore teaching the inner layers by the model itself. Self-distillation allowed the model to outperform other longitudinal models such as [151].

MDITRE ([156]) performed a similar work to phyLoSTM by ordering data phylogenetically and combining both spatial and temporal treatment of the data, while adding visualization with heat maps of the abundance variation over time. They also focused on interpretability by extracting human-readable rules that characterized the evolution of the microbiome. Some of these rules could be sentences like “The average abundance of selected taxa between days 118 and 183 is greater than 7% AND the average slope of selected taxa between days 118 and 190 is greater than 0% per day”. This helps dealing with the problem of how decisions can be taken and justified when relying on black-box models like those found in DL.

The longitudinal paradigm is particularly interesting for retrieving the emergence and progression of a disease over time. Indeed, it is not straightforward to find the causality of a disease in the microbiome using cross-sectional data, and comparing two patients with different diagnosis is also difficult, as the differences between microbiomes may come from very different sources. Studying the same patient at different time points may allow to reduce these sources of discrepancies while increasing the statistical power that could lead to a better understanding of the pathophysiology of the studied disease. To push the idea further, considering the best single-point analysis methods together with LSTM and other longitudinal methods might be key to understanding the most important shifts between healthy and disease states.

#### The reciprocal: predicting microbiome composition

Given that a metagenome can be used to predict phenotype, one can also imagine the other way around. For example, [157] is a k Nearest-Neighbors regression based ML technique which uses species assemblage of a microbiome, ie their absence/presence, to recreate the abundance of each of them without needing complex interaction graphs. In DL, [158] uses phenotype and environmental information to infer the taxonomic composition of the original microbiome without sequencing and binning. Similarly, G2S ([159]) reconstructs the composition of the stool microbiome using information from the dental microbiome: using the abundance table from the dental microbiome diversity, it generates a new abundance table supposed to represent the diversity of the stool microbiome. Finally, to consider temporal data, [160] uses an LSTM to analyze the abundance of a given microbiome at each time step and predict the abundance of the next time-step. This method allows to understand various microbiome dynamics, and can be used to understand the changes in the functions, but also the evolution in metabolite productions.

A recap of methods dealing with phenotype prediction can be found in **Table 2**. A performance comparison can also be found in Table S6.

**Table 2.**
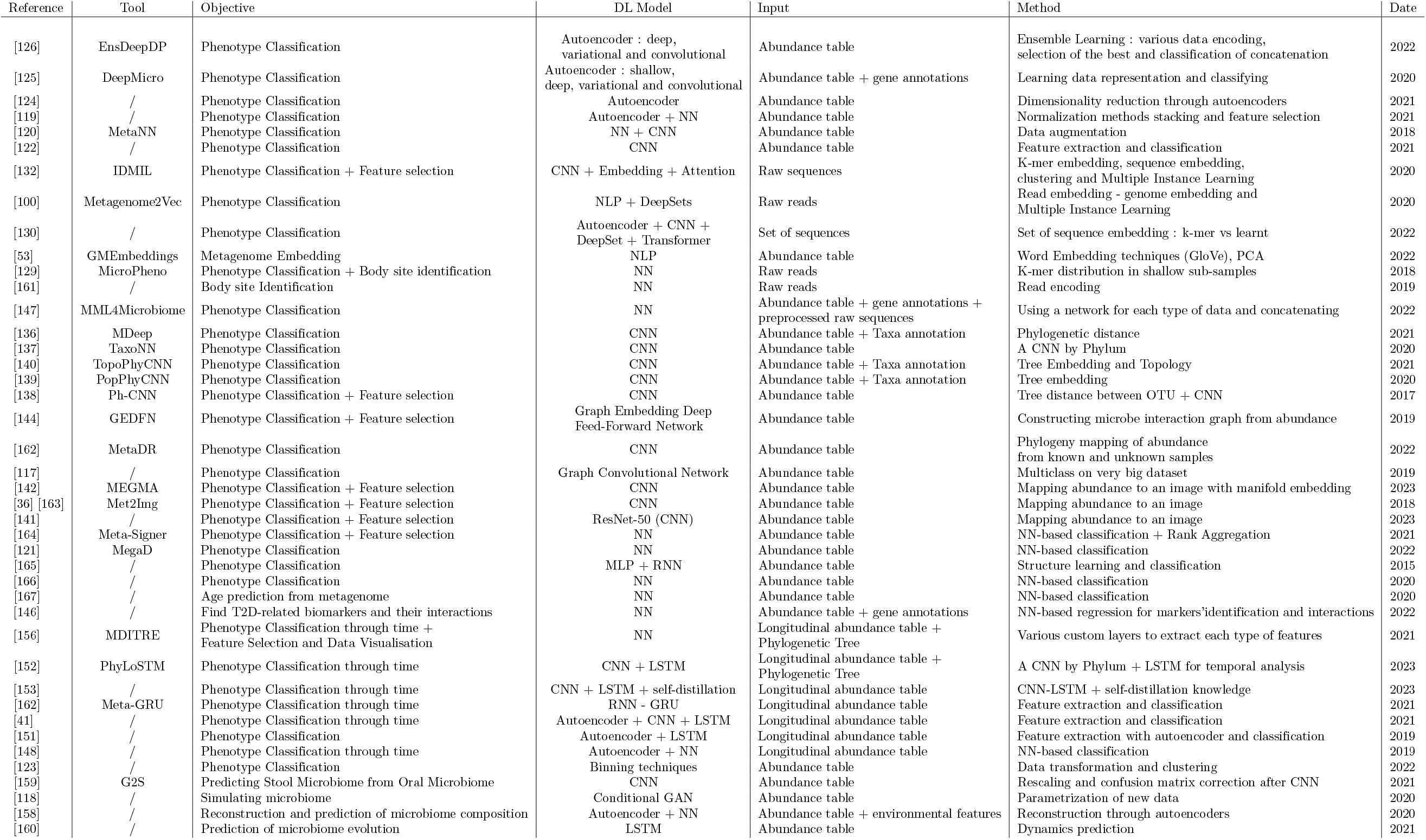
Table of different tools for phenotype prediction. This table summarizes the different tools studied here along with their objective, their input, their model and how they treat information. A table with links to code and dataset and additional information is visible in Supplementary Material

## Discussion

For this metagenomic review, we wanted to focus exclusively on the intersection between the two fields of DL and metagenomics. In need of a reproducible method, we designed a specific search equation. The objective of this equation was to select articles from all other the fields while remaining stringent in order to focus on our theme, as both of the themes composing it present a large litterature. This is why our equation is very specific and searches for words in the title, which can be considered as too stringent. We are aware of this limit, and this is why we decided to enrich our database with connected papers. We are aware that such a choice relies on external tools and leads to choices that can be considered as arbitrary, such as choosing a threshold for the connectivity of articles found via connected papers. We however considered it to be a rich source of data reproducible by anyone. It was important to us having a complete overview of the field, this is why we chose to report together the different steps of metagenomic data analysis and their various scales (sequences, abundance tables, time series, etc…). An overview of the different steps and methods can be seen in **Fig ??**.

Concerning the analysis of various articles, we would like to point out the lack of a solid meta-analysis of DL in metagenomics. It is difficult to compare the performance of the different methods in the literature in a rigorous manner. Indeed, the variety of metagenomic data used, the limited number of samples per study and the recent explosion of many DL methods, do not facilitate their comparison. Most developed methods compare themselves with alignment-based methods or classical machine learning methods like MetaML ([112]), but there is a lack of comparison between DL methods, especially between methods with similar goals but different approaches. Simulated datasets from the CAMI project ([78]) are often used, but even the metrics are difficult to set: good specie classification, quality of bins or differentiation of closely related species. In the case of disease predictions, although datasets are very diverse, the mostly studied diseases form a small group. **Table ??** compiles the different results announced by each articles. These results are obtained on different datasets with various methods and must therefore be treated with caution. However, new arising technologies leave hope for an evergrowing availability with the development of new long read less error prone technologies. Nonetheless, in the case of disease prediction, most studies show a lack in the number of samples, which revolves around a few hundreds of real samples. This is really lower than recommended when using DL, which would rather need thousands of samples for each class. This is a bias that is very important to address with larger accessible complete databases. Before this happens, resampling and data simulation offer an alternative source of data.

Notably, as new powerful DL models are appearing today, we suggest that this meta-analysis will need to include the probably upcoming applications of these models in metagenomics, in particular considering the development of Large Language Models. These models produce impressive results performing many tasks, and their applications to our field will sure be of interest.

These methods, inspired by recent DL technologies like those found in NLP work, such as those mentioned above, are emerging. However, they still need to be further explored, especially using large amounts of data and more powerful models. New powerful Transformer-based models like BERT ([103]) are tested on metagenomic datasets, although adaptations of these approaches to metagenomics are still very recent and the field remains young and very active. Experimental approaches such as BERTax ([105]) and ViBE ([168]) aim to use these powerful models. BERTax, for instance, uses the power of BERT-like models to “learn a representation of the DNA language” and design several taxonomic classification models, at different taxonomic levels (superkingdom, phylum and genus). The ViBE method uses a pre-trained BERT architecture with a reference viral database to identify viruses in the metagenome. Of course, these methods remain challenging because they require very large and representative databases while microbiomes are still composed of many unknown microorganisms. The human gut microbiome is however one of the most studied and described at the genetic and genomic level, with important effort deployed in generating comprehensive catalogs like GTDB ([169]) and GMGC ([25]).

Of course, the quantity of data is of primary importance, but the type of data and the coherence between the pieces of information is just as much of an issue, as it is asked by [147]. As we have seen, classifying a microbiome almost always means using its taxonomic abundance vector, but it must be put into perspective with the question whether microbial communities sorted taxonomically are relevant predictors for these diseases. For a good prediction would be needed communities of microorganisms that are associated in the same way with the studied phenotype. This would mean communities acting positively, negatively or neutrally for a disease in the same way and “quantity”. Taxonomic communities have many advantages, because closely related microbes have a high probability of sharing common behaviors. However, some recent studies have shown that very closely related individuals can behave very differently, sometimes even in opposite ways, despite their taxonomic proximity. This could lead to communities containing microbes both acting positively and negatively, making the community appear neutral. [170] recognizes this problem and proposes a different approach based on guilds. Guilds are based on co-abundance and represent organisms that act in the same direction and therefore evolve together, supposedly in the same dynamics. Questioning the way microorganisms are grouped could be an interesting way to better characterize a metagenome and ultimately improve downstream classification tasks.

Going further than the sole result, one important issue with most of the presented DL approaches is their interpretability. Neural networks are usually black boxes and therefore known to be difficult to understand. However, interpretability is of key importance in the medical field ([47]): understanding how the framework made its decision supports both validation and trust, but also the discovery of novel biomarkers. Many Machine Learning methods are quite useful for interpretability. For example, non-DL methods like MarkerML ([171]) allow the discovery of biomarkers but also the visualization of their interactions. Another non-DL methods, Predomics ([48]), explores the best signatures though very simple models to predict phenotype and allows exploring their features. The high number of transformations and the level of abstraction induced by the layered structure of Neural Networks obscure the way the decision was made. Extracting weights of neurons to assert their importance is one possible solution ([144]), but as the network grows in complexity, it becomes more difficult and unclear. To address this issue, the images created by Met2Img ([36]) are organized using background knowledge such as the ontology of the species. Ablation studies may then be used to identify which parts of the image is most useful to the decision and relate these parts to related species. Besides images, saliency maps can also be calculated to understand which features where mostly used for classification ([172]). Time-evolving methods, by incorporating temporal data, represent a great opportunity in finding new approaches for interpretability, as they permit to extract correlations between changes in features and in phenotype. The rules derived using MDITRE ([156]) are a good step in this direction. The problem remains the fact that microbiome interactions are highly complex and nonlinear, and most of these methods acknowledge the importance of each feature individually, or the comparison of two of them at most, but can hardly give any insight on larger interactions.

## Conclusion

Deep learning has emerged as a promising alternative to traditional bioinformatics approaches in metagenomics in just a few years, for tasks such as binning, sequence prediction, pathogen detection, and phenotype classification. Despite the promising performance of deep learning in metagenomics, a good understanding of the nature of metagenomic data itself remains essential. New sequencing technologies, ever-growing catalogs of species and genes, and studies of microbial interactions may require new approaches to using metagenomic data for disease prediction. Meanwhile, DL and especially powerful transformer-based models such as BERT and GPT, are rapidly advancing and offer significant potential for data analysis in metagenomics, but are still underutilized due to their high computational requirements. These models have a large number of parameters (345 million for BERT and 175 billion for GPT-3), requiring even more data to train effectively. While this data is currently difficult to obtain, its recent and rapid expansion could outpace traditional machine learning algorithms for prediction tasks, paving the way for new models and results. Finally, future work should focus on improving end-to-end analysis of metagenomic data to enable point-of-care applications.

## Acknowledgments

This work was supported by a grant from the French “Agence Nationale de la Recherche” (ANR) for the DeepIntegrOmics project number ANR ANR-21-CE45-0030.

