## Supplementary Material for "Deep learning methods in metagenomics: a review"

October 6, 2023

### 1 Supplementary Data

#### 1.1 Databases

As stated in Material and Methods, we used three databases for our researches : Google Scholar (https://scholar.google.com/), PubMed (https://pubmed.ncbi.nlm.nih.gov/) and IEEE Xplore (https://ieeexplore.ieee.org/Xplore/home.jsp). We later enriched them with Connected papers : https://www.connectedpapers.com/ which is linked with Semantic Scholar Paper Corpus : https://www.semanticscholar.org/paper/Construction-of-the-Literature-Graph-in-Semantic-Amma 649def34f8be52c8b66281af98ae884c09aef38b

All these databases were screened twice, the first time in July 2022 and the second one in July 2023.

#### 1.2 Data Collection

For each article, we collected data on:

- the report: pdf, author, year, and source of publication
- the study: objective, sample characteristics, methods used and features extracted

- the public availability: DOI, url, etc...

- when relevant, the graph from Connected Papers to enrich our dataset

#### 1.3 Filtering

The complete filtering process can be found in our manuscript

#### 1.4 Article Classification

Two screenings were performed, the first one in July 2022 and the second one in July 2023. Tables S7 to S12 and Figures S1 and S3 deal with the first screening, while table S13 deal with the second one. In the first screening, during a second step, we classified manually the articles according to their main objective. This resulted in three main groups: (i) the Taxonomic Classification group with 37 articles, (ii) the Phenotype Prediction group with 49 articles and (iii) a small Miscellaneous group with 4 articles. The first two groups had two articles in common which were omitted (n = 90-2 in common = 88).

This classification was used only for the first screening.

For our synthesis, we used a tag annotation to classify different researches depending on different criteria.

- The objective pursued in the article (binning/sequence identification, disease prediction)

- The input used by the software (raw reads, contigs, abundance table)

- The features extracted from these inputs or external features (abundance, mapping, sequence encoding, k-mer embedding, k-mer distribution, data augmentation, taxonomy, denoising, hashing, co-presence, time series or use of reference database)

- The Deep Learning methods used (classic MLP, autoencoders, CNN, RNN, adversarial networks, NLP, siamese networks and graph embedding)

These tags will help in classifying articles, but have no influence over our selection. Different tags and the number of articles presenting each tag can be found in Tables 9 for goals, 11 for Deep Learning methods, 10 for inputs and 13 for different features used.

#### 1.5 Discussion

##### 1.5.1 Limits in our study

For this metagenomic review, we wanted to focus exclusively on the intersection between the two fields of DL and metagenomics. In need of a reproducible method, we designed a specific search equation. The objective of this equation was to select articles from all other the fields while remaining stringent in order to focus on our theme, as both of the themes composing it present a large litterature. This is why our equation is very specific and searches for words in the title, which can be considered as too stringent. We are aware of this limit, and this is why we decided to enirch our database with connected papers. We are aware that such a choice relies on external tools and leads to choices that can be considered as arbitrary, such as choosing a threshold for the connectivity of articles found via connected papers. We however considered it to be a rich source of data, even though it is less close to the usual systematic review method.

Concerning the analysis of various articles, we would like to point out the lack of a solid meta-analysis of DL in metagenomics. This is due to several reasons we detail further in this section. However, as new powerful DL models, are appearing today, we suggest that this meta-analysis will need to include the probably upcoming applications of these models in metagenomics. These models produce impressive results performing many tasks, and their applications to our field will sure be of interest.

##### 1.5.2 Datasets discussion

Although metagenomic data is getting increasingly easy to collect and analyze thanks to rapidly evolving new technologies, there is a lack of reference datasets on which could be compared the different tools developed for analysis. This is especially true in phenotype prediction. At the sequence level, simulation technologies such as CAMISIM make it easier to generate toy samples and therefore having a better comparison, but there are still limits. Moreover, as most of the microbes found in human microbiome are still unknown, catalogs are still widely incomplete, although they are quickly developing. All these limitations make it difficult to compare the newly developed tools between one another. As DL massive deployment is still recent, and because of the most methods do not compare themselves to other DL methods, but rather to classic or ML methods, which makes it difficult to compare their performances.

##### 1.5.3 State and Future of the field

As said, DL is quite a recent technology. However, its rapidly evolving methods have been quickly taken into account by bioinformaticians, and most metagenomic issues can nw be addressed through DL. All classic models found their use, be it classic MLP, CNN, LSTM or, more recently transformers. If some architectures seem well-suited for some types of tasks, for example CNN when studying abundance table and phylogeny or LSTM for longitudinal data, sometimes combined with CNN, many different have been tested. Of course, applications in metagenomics can take a little time to keep up with the rapidly advancing field of DL. Therefore, applications of most recent groundbreaking transformer models are still rare and developing. Attention-based models or even Transformers like BERT are already used, but many tasks seem to fit to their advantages.

Therefore, in our time of flourishing Large Language Models, it would be logical to see the development of such models specifically applied to metagenomics. These models could learn the structure of DNA and therefore show performances that could be as impressive as they already are in many other fields. Concerning the datasets, new arising technologies leave hope for an evergrowing availability with the development of new long read less error prone technologies.

#### 1.6 Public Availability

The availability of the articles retrieved are visible in Table S8.

#### 1.7 Data Availability

All publicly available or open-access articles are described in Table "Table of all studies". Notebook and codes used to sort and filter articles and enrich via Connected Papers can be found on `https://github.com/CorvusVaine/analyzing_connected_papers_articles.git`

#### 1.8 Authors

This section deals with authors participation in screening. During the screening phase, the search equation was designed by Gaspar Roy and validated by Edi Prifti. The enrichment method was jointly designed and discussed by Gaspar Roy, Edi Prifti and Jean-Daniel Zucker. Both these methods were discussed between authors in order to keep a reproducible screening method being exhaustive while stringent enough. The enrichment aimed to leave no important article behind in order to avoid strong biases that could have been induced by the research equation. Finally, the filtering was performed to make sure this enrichment did not create new biases by adding articles related to only one of the themes but not the other one. The screening and synthesis was made by Gaspar Roy under the direction and review of Edi Prifti, Eugeni Belda and Jean-Daniel Zucker.

#### 1.9 Table Preparation

Tables 1 and 7 have been prepared to summarize the various analyses cited in the review performing on metagenomic data. They were separated in two levels depending both on the data used and the objective pursued. The separation is therefore between work at sequence level and work at metagenomic profile level and hence between predicting the nature or origin of a sequence, and predicting a phenotype or visalizing the function of a microbiome. Inside each table, we then sorted the articles by their more accurate objective : classifying if a sequence is of a certain type, taxonomic classification, binning, or on the other side phenotype prediction, temporary data, etc.. We tried the most in this classification to sort methods depending on their DL architecture and then, finally, depending on the date they were released. These tables act as an inventory of all methods, but do not ambition to say much on their performances. We tried to do so in other tables. As we shall say in the **Discussion**, comparison between methods is still a quite open field, and is therefore not an easy task. We tried to give an idea by citing in Table 8 the different results claimed by each method depending on the predicted disease. But if there is a limited number of diseases that can be assessed from microbiome data, many datasets exist, which can be a source of biases, as many parameters can influence one's health including ethnicity or alimentation. In the same idea, many different datasets can be used for sequence analysis, as seen in table 2. However, the possibility to simulate reads with tools such as CAMISIM opens new perspectives for comparison. We took the time to make Tables 3,4,5 and 6. Indeed, VAMB set a standard in DL Binning, and most recent advances compared themselves to this tool and not only classic bioinformatics tools or Machine Learning methods. Therefore, it appeared to us of interest to the user to have these information. It is noteworthy that the comparison to VAMB was always made by the authors of the latter method.

#### 2 Details on our first screening

This section mostly contains details, figures, tables and statistics concerning the research method of articles. Connected Papers can be found at `https://www.connectedpapers.com/` Two screenings were performed, the first one in July 2022 and the second one in July 2023. Tables S1 to S6 and Figures S1 and S3 deal with the first screening, while table S7 deal with the second one. In the first screening, during a second step, we classified manually the articles according to their main objective. This resulted in three main groups: (i) the Taxonomic Classification group with 37 articles, (ii) the Phenotype Prediction group with 49 articles and (iii) a small Miscellaneous group with 4 articles. The first two groups had two articles in common which were omitted (n = 90-2 in common = 88). We then generated a graph for each of the first two groups (respectively named $G_{tc}$ and $G_{pp}$), the miscellaneous group was discarded as it contained irrelevant articles. The connectivity distribution (i.e. the number of links for each node) was higher in the Phenotype Prediction group compared to Sequence Classification, with a large number of already selected articles pointing to each other. It is possible to see different graphs of articles for different co-citation connectivity thresholds along with the distribution of articles in **Figure S3**. We decided to add to our dataset the articles with a co-citation connectivity $> 4$ for both graphs $G_{tc}$ and $G_{pp}$. This filtering allowed us to recover 21 and 37 new articles, respectively. Using all articles together to form a graph (called $G_{all}$), we recovered 10 more articles with the same threshold. We chose this threshold because it allowed us to reject as few articles as possible while not adding more articles than the original database size. For example, given a total of 114 articles, a threshold of three identifies 147 additional articles, more than doubling the dataset of articles. We therefore chose a threshold of four, which added 68 previously unseen articles. A majority of the newly discovered articles deal with the Phenotype Prediction group rather than Sequence Classification : 37 were found with $G_{pp}$, of which 12 (36%) were retained after filtering, 21 were found with $G_{tc}$, 2 after filtering (9.5%) and 68 were found considering $G_{all}$ , with 17 conserved(25%). After filtering, these 17 supplementary articles are kept and added to the initial corpus for further analyses. No thematic sort was preformed for the second screening.

### 3 Supplementary Tables and Figures

#### 3.1 Performance Tables

This section contains performance tables on different types of methods. It is to be noted that the performances shown here are giver by articles studied and not by an independant analysis.

Tables S1, S2, S3, S4 and S5 deal with sequence-based methods.

| Article | Simulated Short Reads | Simulated Long Reads | Real Short Reads | Real Long Reads |
|---|---|---|---|---|
| [1] | F-measure : 0.8716 | F-measure : 0.9595 | F-measure : 0.701 | / |
| [2] | / | / | Accuracy on genus-level : 0.761 | / |
| [3] | / | / | F1 on species : 0.9894 | / |
| [4] | / | / | V-Measure : 0.932 | / |
| [5] | / | / | Species : Precision : 80 and Recall : 0.891 | / |
| [6] | / | / | Genus Accuracy : 0.913 | / |
| [7] | / | / | Genus Precision : 0.969 and Recall : 0.866 | / |
| [8] | / | / | Clustering V-measure : 0.932 | / |
| [9] | / | / | Species-level F1 : 0.907 | / |
| [10] | / | / | / | Species Precision : 0.973 and Recall : 0.3305 |
| [11] | / | / | Species-level on 100 genomes : 0.8864 | / |
| [12] | / | / | Species-level Accuracy : 0.739 | / |
| [13] | / | Accuracy : 0.783 | / | / |
| [14] | F1 : 0.6589 | / | / | / |
| [15] | / | / | Phylum-level Accuracy : 0.601 | / |
| [16] | Accuracy : 0.8795 | Accuracy : 0.9844 | / | / |

Table S 1: **Table listing different DL-based methods as well as their performance in taxonomic classification.** Please note that the results are found on each model's own dataset, not on a centralized dataset, using different metrics and data of various complexity, and can therefore not easily be compared.

| Method | Airways | GI | Oral | Skin | Urog |
|---|---|---|---|---|---|
| VAMB | 143 | 180 | 142 | 284 | 131 |
| CLMB | 144 | 201 | 163 | 253 | 155 |

Table S 2: **Table of comparison of performances in High-Quality genome recovery.** This table compares the number of genomes retrieved by VAMB ([17] and CLMB ([18]) as described in the latter article. This comparison takes place over real datasets from Airways, Gastrointestinal, Oral, Skin and Urogenital microbiome.

| Method | Simulated Skin | Simulated Oral | Real Human Gut | Real Dog Gut | Real Ocean | Real Soil |
|---|---|---|---|---|---|---|
| VAMB | 63 | 88 | 344 | 97 | 233 | 29 |
| SemiBin | 75 | 108 | 368 | 100 | 314 | 81 |

Table S 3: **Table of comparison of performances in High-Quality distinct species with multi-sample binning.** This table compares the number of high-quality distinct species returned with multi-sample binning by VAMB ([17] and SemiBin ([19]) as described in the latter article. This comparison takes place over simulated CAMI datasets of Skin and Oral microbiome, as well as real datasets from Human gut, dog gut, soil and ocean microbiome.

| Method | Airways | Gastro-intestinal | Oral | Skin | Urogenital | Total |
|---|---|---|---|---|---|---|
| VAMB | 63 | 82 | 124 | 72 | 78 | 440 |
| AAMB | 74 | 98 | 118 | 90 | 70 | 472 |
| AVAMB | 82 | 103 | 138 | 104 | 83 | 532 |
| MetaBAT2 | 36 | 76 | 68 | 62 | 66 | 309 |
| SemiBin | 96 | 140 | 159 | 138 | 112 | 645 |

Table S 4: **Table of comparison of performances in Near-Complete Genomes reconstructed from the CAMI2 datasets.** This table compares the number of near-complete genomes reconstructed by VAMB ([17]), MetaBAT2 ([20]), SemiBin ([19]), AAMB and AVAMB ([21]) as described in the preprint of the latter method. This comparison takes place over simulated CAMI datasets of Airways, Gastro-intestinal, Oral, Skin and Urogenital microbiome.

| Method | Aale | Mari | Damh | Hjor | Hade | Viby | Total |
|---|---|---|---|---|---|---|---|
| MetaBAT2 | 53 | 41 | 50 | 28 | 51 | 30 | 309 |
| VAMB | 42 | 37.3 | 41.3 | 22 | 47.3 | 19 | 208.9 |
| VAE+E+SCG | 60.4 | 47.4 | 49.8 | 27.8 | 52.4 | 29 | 266.2 |

Table S 5: **Table of comparison of performances in High-quality bins.** This table compares the number of high-quality bins reconstructed by VAMB ([17]), MetaBAT2 ([20]) and VAE+E+SCG ([22]) as described in the latter's article. This comparison takes place over Wastewater Treatment Plant dasasets.

Table S6 above is also a performance comparison, dealing this time with disease prediction from abundance tables.

| Article | CRC | IBD | CIR | OBE | T2D | W2D | CrD | UC | Year | Tool |
|---|---|---|---|---|---|---|---|---|---|---|
| [23] | / | / | / | / | Mean AUC : 0.811 | / | / | / | 2022 | / |
| [24] | 0.834 | 0.844 | 0.949 | 0.677 | 0.776 | 0.786 | / | / | 2022 | / |
| [25] | / | 0.81 | / | / | / | / | / | / | 2022 | GMEmbeddings |
| [26] | 0.894 | 0.941 | 0.911 | 0.714 | 0.771 | 0.860 | / | / | 2022 | EnsDeepDP |
| [27] | 0.803 | 0.955 | 0.940 | 0.659 | 0.763 | 0.899 | / | / | 2020 | DeepMicro |
| [28] | / | 0.89 | / | / | / | / | / | / | 2019 | MetaNN |
| [29] | 0.818 | 0.863 | 0.862 | 0.656 | 0.564 | 0.704 | / | / | 2021 | CNN1D |
| [30] | 0.857 to 0.987 | / | / | / | / | / | / | / | 2021 | / |
| [31] | 0.895 | 0.882 | 0.951 | 0.793 | 0.816 | / | / | / | 2020 | IDMIL |
| [32] | 0.81 | / | 0.83 | / | / | / | / | / | 2020 | Metagenome2Vec |
| [33] | / | / | / | / | / | / | 0.884 | / | 2022 | / |
| [34] | / | / | / | / | / | / | 0.76 | / | 2018 | MicroPheno |
| [35] | 0.988 | 0.991 | 0.886 | / | 0.735 | / | / | / | 2022 | MML4Microbiome |
| [36] | / | / | 0.946 | 0.666 | 0.69 | / | / | / | 2020 | PopPhyCNN |
| [37] | / | / | / | / | / | / | 0.926 | 0.946 | 2018 | Ph-CNN |
| [38] | / | / | 0.938 | / | 0.762 | / | / | / | 2020 | TaxoNN |
| [39] | / | 0.843 | / | / | / | / | / | / | 2019 | GEDFN |
| [40] | 0.9063 | / | 0.9535 | / | 0.7890 to 0.8131 | / | / | / | 2022 | EPCNN |
| [41] | Top 1 : 0.36 / Top 5 : .84 | Top 1 : 0.4 / Top 5 : 0.94 | / | / | / | / | / | / | 2019 | / |
| [42] | F1 : 0.549 | AUC : 0.940 | 0.949 | 0.642 | 0.740 | / | / | / | 2023 | MEGMA |
| [43] [44] | 0.820 | / | 0.926 | 0.696 | / | 0.749 | / | / | 2020 | Met2Img |
| [45] | / | / | / | / | 0.96 | / | / | / | 2023 | / |
| [46] | / | / | 0.833 | / | 0.7 | / | / | / | 2022 | MegaD |
| [47] | / | / | / | / | / | / | / | 0.889 | 2023 | / |

Table S 6: **Table of different tools' performances in predicting various diseases.** The given score is ROC AUC, the diseases are Colorectal Cancer (CRC), Inflammatory Bowel Disease (IBD), Cirrhosis (CIR), Obesity (OBE), Type 2 Diabetes (T2D and W2D), Crohn Disease (CrD) and Ulcerative Colitis (UC). The results are found on each model's own dataset, not on a centralized dataset, which limits their comparability.

#### 3.2 Figures

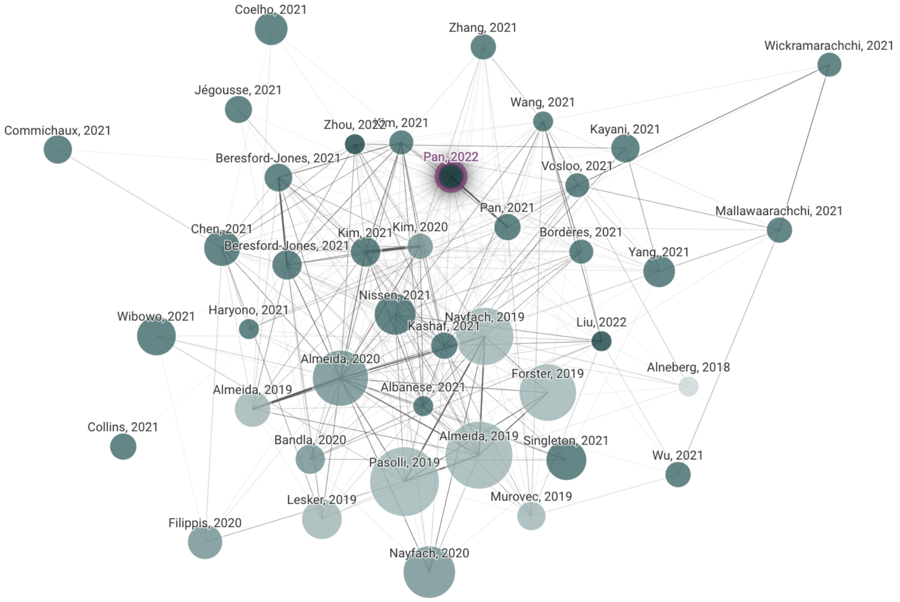


Fig S 1: This is an example of a graph generated using Connected Papers. It has been generated from article "A deep siamese neural network improves metagenome-assembled genomes in microbiome datasets across different environments" ([19]). The darker the color is, the more recent the article, and the bigger the circle is, the most citations it has.

#### 3.3 First Screening Tables

#### 3.4 Connectivity of newly found papers

This paragraph concerns the first screening, happening in July 2022. The new relevant articles we selected with the Connected Papers approach had a co-citation connectivity greater than 4. However, we can question whether there is a difference between retained and filtered articles in terms of co-citation connectivity. Therefore, we will know if articles who passed the abstract filtering

| Number of links | 1 | 2 | 3 | 4 | 5 | 6 | 7 | 8 | 9 | 10 | 11 | 12 | 13 | 14 | 15 | 16 | 17 | 18 | 19 | 20 | 21 | 22 | 23 |
|---|---|---|---|---|---|---|---|---|---|---|---|---|---|---|---|---|---|---|---|---|---|---|---|
| Sequence classification Total | 1029 | 210 | 83 | 42 | 28 | 13 | 6 | 6 | 6 | 6 | NA | NA | NA | NA | NA | NA | NA | NA | NA | NA | NA | NA | NA |
| Sequence classification New | 992 | 179 | 58 | **21** | 12 | 2 | 1 | 1 | 1 | 1 | NA | NA | NA | NA | NA | NA | NA | NA | NA | NA | NA | NA | NA |
| Phenotype prediction Total | 1069 | 239 | 116 | 72 | 63 | 59 | 54 | 50 | 49 | 48 | 47 | 46 | 42 | 42 | 42 | 39 | 34 | 31 | 30 | 30 | 21 | NA | NA |
| Phenotype prediction New | 1022 | 197 | 79 | **37** | 31 | 27 | 23 | 19 | 18 | 17 | 16 | 15 | 12 | 12 | 12 | 10 | 6 | 5 | 4 | 4 | 1 | NA | NA |
| All papers Total | 2065 | 512 | 215 | 129 | 98 | 85 | 67 | 61 | 59 | 58 | 52 | 51 | 49 | 47 | 47 | 46 | 38 | 38 | 34 | 34 | 28 | 28 | 23 |
| All papers New | 1979 | 433 | 147 | **68** | 46 | 34 | 26 | 21 | 19 | 18 | 17 | 16 | 15 | 13 | 13 | 12 | 6 | 6 | 4 | 4 | 1 | 1 | 0 |

Table S 7: Distribution of new articles discovered by number of original articles pointing to them. 0 means there were articles with such connectivity but none of them were new

| Origin | Sequence Classification Only | Phenotype Prediction Only | All articles |
|---|---|---|---|
| Raw | 21 | 37 | 68 |
| Filtered | 2 | 12 | 17 |

Table S 8: Distribution of new articles and kept new articles by origin

| Taxonomy/Sequence Identification | Phenotype Classification | Miscellaneous |
|---|---|---|
| 58 | 39 | 4 |

Table S 9: Distribution of articles by goal of the article.

| raw reads | contigs | abundance table |
|---|---|---|
| 36 | 10 | 48 |

Table S 10: Distribution of articles by type of input.

| MLP | Autoencoders | CNN | RNN | Adversarial | NLP | Siamese | Graph embedding | ML Methods |
|---|---|---|---|---|---|---|---|---|
| 28 | 17 | 31 | 8 | 3 | 20 | 3 | 2 | 20 |

Table S 11: Distribution of articles by Deep Learning methods.

| Abundance | Mapping | sequence encoding | Image encoding | k-mer embedding | k-mer distribution | data augmentation | taxonomy | denoising | hashing | Co-presence | Time series | Reference database |
|---|---|---|---|---|---|---|---|---|---|---|---|---|
| 46 | 10 | 14 | 5 | 8 | 16 | 3 | 14 | 3 | 4 | 7 | 8 | 15 |

Table S 12: Distribution of articles by features used.

| Number of links | 0 | 1 | 2 | 3 | 4 | 5 | 6 | 7 | 8 | 9 | 10 | 11 | 12 | 13 | 14 | 15 | 16 | 17 | 18 | 19 | 20 | 21 | 22 | 23 | 24 | 25 | 26 | 27 | 28 | 29 | 30 | 31 | 32 | 33 | 34 |
|---|---|---|---|---|---|---|---|---|---|---|---|---|---|---|---|---|---|---|---|---|---|---|---|---|---|---|---|---|---|---|---|---|---|---|---|
| Number of new papers | 2443 | 2443 | 638 | 260 | **130** | 94 | 73 | 55 | 40 | 32 | 27 | 23 | 22 | 20 | 19 | 17 | 14 | 13 | 12 | 12 | 9 | 6 | 5 | 5 | 5 | 4 | 4 | 3 | 3 | 3 | 2 | 2 | 1 | 1 | 1 |
| Total of papers | 2550 | 2550 | 736 | 343 | **201** | 162 | 136 | 118 | 100 | 92 | 84 | 72 | 68 | 66 | 58 | 56 | 53 | 52 | 50 | 50 | 47 | 42 | 39 | 39 | 39 | 37 | 37 | 36 | 34 | 34 | 32 | 32 | 30 | 30 | 30 |

Table S 13: Distribution of new articles discovered by number of original articles pointing to them in second screening. 0 means there were articles with such connectivity but none of them were new

| Title | DOI | Publicly available | Date |
|---|---|---|---|
| Phylogenetic convolutional neural networks in metagenomics | 10.1186/s12859-018-2033-5 | Yes | March 8, 2018 |
| Using convolutional neural networks to explore the microbiome | 10.1109/EMBC.2017.8037799 | Yes | July, 2017 |
| Improved metagenome binning and assembly using deep variational autoencoders | 10.1038/s41587-020-00777-4 | Yes | January 4, 2021 |
| Identifying viruses from metagenomic data using deep learning | 10.1007/s40484-019-0187-4 | Yes | March 8, 2020 |
| CNN-MGP: Convolutional Neural Networks for Metagenomics Gene Prediction | 10.1007/s12539-018-0313-4 | Yes | December 11, 2021 |
| DeepMicrobes: taxonomic classification for metagenomics with deep learning | 10.1093/nargab/lqaa009 | Yes | February 19, 2020 |
| DeepARG: a deep learning approach for predicting antibiotic resistance genes from metagenomic data | 10.1186/s40168-018-0401-z | Yes | February 1, 2021 |
| PopPhy-CNN: A Phylogenetic Tree Embedded Architecture for Convolutional Neural Networks to Predict Host Phenotype From Metagenomic Data | 10.1109/JBHI.2020.2993761 | Yes | January 31, 2018 |
| Deep learning models for bacteria taxonomic classification of metagenomic data | 10.1186/s12859-018-2182-6 | Yes | July 9, 2018 |
| A deep siamese neural network improves metagenome-assembled genomes in microbiome datasets across different environments | 10.1038/s41467-022-29843-y | Yes | April 28, 2022 |
| Interpretable and accurate prediction models for metagenomics data | 10.1093/gigascience/giaa010 | Yes | March 1, 2021 |
| Identification of antimicrobial peptides from the human gut microbiome using deep learning | 10.1038/s41587-022-01226-0 | No | March 3, 2022 |
| MetaPheno: A critical evaluation of deep learning and machine learning in metagenome-based disease prediction | 10.1016/j.ymeth.2019.03.003 | Yes | August 15, 2019 |
| Interpretable Machine Learning Framework Reveals Robust Gut Microbiome Features Associated With Type 2 Diabetes | 10.2337/dc20-1536 | Yes | December 7, 2020 |
| Enhanced metagenomic deep learning for disease prediction and consistent signature recognition by restructured microbiome 2D representations | 10.1016/j.patter.2022.100658 | Yes | January 13, 2023 |
| Human Gut Microbiome Aging Clock Based on Taxonomic Profiling and Deep Learning | 10.1016/j.isci.2020.101199 | Yes | May 23, 2020 |
| phyLoSTM: a novel deep learning model on disease prediction from longitudinal microbiome data | 10.1093/bioinformatics/btab482 | Yes | July 2, 2021 |
| Multimodal deep learning applied to classify healthy and disease states of human microbiome | 10.1038/s41598-022-04773-3 | Yes | January 17, 2022 |
| A novel deep learning method for predictive modeling of microbiome data | 10.1093/bib/bbaa073 | Yes | May 20, 2021 |
| Deep Learning for Metagenomic Data: using 2D Embeddings and Convolutional Neural Networks | 10.48550/arXiv.1712.00244 | Yes | December 1, 2017 |
| Towards a metagenomics machine learning interpretable model for understanding the transition from adenoma to colorectal cancer | 10.1038/s41598-021-04182-y | Yes | January 10, 2022 |
| Machine learning and deep learning applications in microbiome research | 10.1038/s43705-022-00182-9 | Yes | October 6, 2022 |
| Interpretable machine learning framework reveals microbiome features of oral disease | 10.1101/2020.04.05.024984 | Yes | December, 2022 |
| Predicting microbiome compositions from species assemblages through deep learning | 10.1002/imt2.3 | Yes | March 1, 2022 |
| CHEER: HierarCHical taxonomic classification for viral mEtagEnomic data via deep leaRning | 10.1016/j.ymeth.2020.05.018 | Yes | May 23, 2020 |
| PPR-Meta: a tool for identifying phages and plasmids from metagenomic fragments using deep learning | 10.1093/gigascience/giz066 | Yes | June 20, 2019 |
| MarkerML – Marker Feature Identification in Metagenomic Datasets Using Interpretable Machine Learning | 10.1016/j.jmb.2022.167589 | No | June 15, 2022 |
| Taxonomic classification of metagenomic sequences from Relative Abundance Index profiles using deep learning | 10.1016/j.bspc.2021.102539 | No | May, 2021 |
| CoCoNet: an efficient deep learning tool for viral metagenome binning | 10.1093/bioinformatics/btab213 | Yes | April 5, 2021 |
| Disease Classification in Metagenomics with 2D Embeddings and Deep Learning | 10.48550/arXiv.1806.09046 | Yes | June 23, 2018 |
| MetaVelvet-DL: a MetaVelvet deep learning extension for de novo metagenome assembly | 10.1186/s12859-020-03737-6 | Yes | June 2, 2021 |
| Deep in the Bowel: Highly Interpretable Neural Encoder-Decoder Networks Predict Gut Metabolites from Gut Microbiome | 10.1186/s12864-020-6652-7 | Yes | July 20, 2020 |
| Machine Learning and Deep Learning Applications in Metagenomic Taxonomy and Functional Annotation | 10.3389/fmicb.2022.811495 | Yes | March 14, 2022 |
| Human microbiome aging clocks based on deep learning and tandem of permutation feature importance and accumulated local effects | 10.1101/507780 | Yes | December 28, 2018 |
| Disease Prediction Using Synthetic Image Representations of Metagenomic Data and Convolutional Neural Networks | 10.1109/RIVF.2019.8713670 | No | May 16, 2019 |
| PopPhy-CNN: A Phylogenetic Tree Embedded Architecture for Convolution Neural Networks for Metagenomic Data | 10.1109/JBHI.2020.2993761 | Yes | May 11, 2020 |
| DeepToA: an ensemble deep-learning approach to predicting the theater of activity of a microbiome | 10.1093/bioinformatics/btac584 | Yes | August 27, 2022 |
| Gene Prediction in Metagenomic Fragments with Deep Learning | 10.1155/2017/4740354 | Yes | November 8, 2017 |
| Feature Extension of Gut Microbiome Data for Deep Neural Network-Based Colorectal Cancer Classification | 10.1109/ACCESS.2021.3050838 | Yes | January 11, 2021 |
| A novel thermophilic chitinase directly mined from the marine metagenome using the deep learning tool Preoptem | 10.1186/s40643-022-00543-1 | Yes | May 16, 2022 |
| Human disease prediction from microbiome data by multiple feature fusion and deep learning | 10.1016/j.isci.2022.104081 | Yes | April 15, 2022 |
| MDITRE: Scalable and Interpretable Machine Learning for Predicting Host Status from Temporal Microbiome Dynamics | 10.1128/msystems.00132-22 | Yes | September 7, 2022 |
| Novel taxonomy-independent deep learning microbiome approach allows for accurate classification of different forensically relevant human epithelial materials | 10.1016/j.fsigen.2019.03.015 | Yes | July, 2019 |
| Application of Deep Learning in Microbiome | 10.2991/jaims.d.201028.001 | Yes | January, 2020 |
| EnsDeepDP: An Ensemble Deep Learning Approach for Disease Prediction Through Metagenomics | 10.1109/TCBB.2022.3201295 | No | August 24, 2022 |
| IDMIL: an alignment-free Interpretable Deep Multiple Instance Learning (MIL) for predicting disease from whole-metagenomic data | 10.1093/bioinformatics/btaa477 | Yes | July 13, 2020 |
| Deep Learning Tools for Human Microbiome Big Data | 10.1007/978-3-319-62521-8_21 | No | August, 2018 |
| Utilizing longitudinal microbiome taxonomic profiles to predict food allergy via long short-term memory networks | 10.1371/journal.pcbi.1006693 | Yes | February 4, 2019 |
| DLmeta: a deep learning method for metagenomic identification | 10.1109/BIBM55620.2022.9995231 | No | December 2022 |
| Dimensionality Reduction for Cluster Identification in Metagenomics using Autoencoders | 10.1109/ICTer51097.2020.9325447 | Yes | November 2020 |
| Stacking and Chaining of Normalization Methods in Deep Learning-Based Classification of Colorectal Cancer Using Gut Microbiome Data | 10.1109/ACCESS.2021.3094529 | Yes | July 2021 |
| Metagenome-Based Disease Classification with Deep Learning and Visualizations Based on Self-organizing Maps | 10.1007/978-3-030-35653-8_20 | No | November 2019 |
| Ontology-aware neural network: a general framework for pattern mining from microbiome data | 10.1093/bib/bbac005 | Yes | March, 2022 |
| Predicting Host Phenotype Based on Gut Microbiome Using a Convolutional Neural Network Approach | 10.1007/978-1-0716-0826-5_12 | No | August 18, 2020 |
| TopoPhy-CNN: Integrating Topological Information of Phylogenetic Tree for Host Phenotype Prediction From Metagenomic Data | 10.1109/BIBM52615.2021.9669509 | No | 2021 |
| NLP-MeTaxa: A Natural Language Processing Approach for Metagenomic Taxonomic Binning Based on Deep Learning | 10.2174/1574893616666210621101150 | Yes | June, 2021 |
| Prediction of microbial communities for urban metagenomics using neural network approach | 10.1186/s40246-019-0224-4 | Yes | October 22, 2019 |
| GraphKKE: graph Kernel Koopman embedding for human microbiome analysis | 10.1007/s41109-020-00339-2 | Yes | December 1, 2020 |
| Understanding microbiome dynamics via interpretable graph representation learning | 10.1038/s41598-023-29098-7 | Yes | February 4, 2023 |
| MegaD: Deep Learning for Rapid and Accurate Disease Status Prediction of Metagenomic Samples | 10.3390/life12050669 | Yes | April 30, 2022 |
| G2S: A New Deep Learning Tool for Predicting Stool Microbiome Structure From Oral Microbiome Data | 10.3389/fgene.2021.644516 | Yes | April 9, 2021 |
| Classification of Microbiome Data from Type 2 Diabetes Mellitus Individuals with Deep Learning Image Recognition | 10.3390/bdcc7010051 | Yes | March 17, 2023 |
| Disease Prediction Using Metagenomic Data Visualizations Based on Manifold Learning and Convolutional Neural Network | 10.1007/978-3-030-35653-8_9 | No | November 20, 2019 |
| A self-knowledge distillation-driven CNN-LSTM model for predicting disease outcomes using longitudinal microbiome data | 10.1093/bioadv/vbad059 | Yes | May 18, 2023 |
| PlasGUN: gene prediction in plasmid metagenomic short reads using deep learning | 10.1093/bioinformatics/btaa103 | Yes | May, 2020 |
| Deep Learning Enables Design of Multifunctional Synthetic Human Gut Microbiome Dynamics | 10.1101/2021.09.27.461983 | Yes | September 28, 2021 |
| ResMiCo: Increasing the quality of metagenome-assembled genomes with deep learning | 10.1371/journal.pcbi.1011001 | Yes | May 1, 2023 |
| TAMPA: interpretable analysis and visualization of metagenomics-based taxon abundance profiles | 10.1093/gigascience/giad008 | Yes | February 28, 2023 |
| Deep Learning Encoding for Rapid Sequence Identification on Microbiome Data | 10.3389/fbinf.2022.871256 | Yes | June 24, 2022 |
| Using Autoencoders for Predicting Latent Microbiome Community Shifts Responding to Dietary Changes | 10.1109/BIBM47256.2019.8983124 | Yes | November, 2019 |
| Early Identification of Fungal and Mycobacterium Infections in Pulmonary Granulomas Using Metagenomic Next-Generation Sequencing on Formalin fixation and paraffin embedding tissue | 10.1080/14737159.2022.2052046 | No | April 22, 2022 |
| Life Language Processing: Deep Learning-based Language-agnostic Processing of Proteomics, Genomics/Metagenomics, and Human Languages | / | Yes | 2019 |
| SemiBin: Incorporating information from reference genomes with semi-supervised deep learning leads to better metagenomic assembled genomes (MAGs) | 10.1101/2021.08.16.456517 | Yes | August 16, 2021 |
| Deep learning and host variable embedding augment microbiome-based simultaneous detection of multiple diseases | 10.1101/2023.05.16.541058 | Yes | May 18, 2023 |
| NLP-based classification of software tools for metagenomics sequencing data analysis into EDAM semantic annotation | 10.48550/arXiv.2210.00831 | Yes | September 30, 2022 |
| Reject and Cascade Classifier with Subgroup Discovery for Interpretable Metagenomic Signatures | 10.1007/978-3-030-93736-2_5 | Yes | February 17, 2022 |
| MT-MAG: Accurate and interpretable machine learning for complete or partial taxonomic assignments of metagenome-assembled genomes | 10.1101/2022.01.12.475159 | Yes | June 21, 2023 |
| GMEmbeddings: An R Package to Apply Embedding Techniques to Microbiome Data | 10.3389/fbinf.2022.828703 | Yes | April 26, 2022 |
| Improving Disease Prediction using Shallow Convolutional Neural Networks on Metagenomic Data Visualizations based on Mean-Shift Clustering Algorithm | 10.14569/IJACSA.2020.0110607 | Yes | January, 2020 |
| DeepMicro: deep representation learning for disease prediction based on microbiome data | 10.1038/s41598-020-63159-5 | Yes | April 7, 2020 |
| MetaMLP: A Fast Word Embedding Based Classifier to Profile Target Gene Databases in Metagenomic Samples | 10.1089/cmb.2021.0273 | No | November 28, 2021 |
| A Deep Learning Approach to Predict Health Status Using Microbiome Profiling | 10.1109/ICTer58063.2022.10024102 | No | November 30, 2022 |
| Adversarial and variational autoencoders improve metagenomic binning | 10.1101/2023.02.27.527078 | Yes | February 27, 2023 |
| Interpretable machine learning analysis of functional metagenomic profiles improves colorectal cancer prediction and reveals basic molecular mechanisms | 10.21203/rs.2.21634/v1 | Yes | January, 2020 |
| Neural network-based taxonomic clustering for metagenomics | 10.1109/IJCNN.2010.5596644 | Yes | July, 2010 |
| Embedding the de Bruijn graph, and applications to metagenomics | 10.1101/2020.03.06.980979 | Yes | March 8, 2020 |
| Interpretable Machine Learning Algorithm Reveals Novel Gut Microbiome Features in Predicting Type 2 Diabetes | 10.1093/cdn/nzaa062_016 | Yes | June, 2020 |
| Metagenomic Binning using Connectivity-constrained Variational Autoencoders | id8OSRGeew | Yes | April 28, 2023 |
| Deep Learning Approach for Pathogen Detection Through Shotgun Metagenomics Sequence Classification | 10.1007/978-3-030-21642-9_4 | No | May 30, 2019 |
| Feature Selection Based on a Shallow Convolutional Neural Network and Saliency Maps on Metagenomic Dat | 10.1007/978-981-33-6385-4_10 | No | January, 2021 |
| Metagenomic Sequence Classification based on One-Dimensional Convolutional Neural Network | 10.1145/3581807.3581835 | Yes | 2022 |
| A K-mer based Multi Convolutional Neural Network Classifier of Low-Ranking Taxonomic Bins from Metagenome | 335540811 | Yes | September, 2019 |
| Feature Selection Using Local Interpretable Model-Agnostic Explanations on Metagenomic Data | 10.1007/978-3-030-76620-7_6 | No | June, 2021 |
| DephageTP: a convolutional neural network framework for identifying phage-specific proteins from metagenomic sequencing data | 10.7717/peerj.13404 | Yes | June 8, 2022 |
| A neural network-based framework to understand the type 2 diabetes-related alteration of the human gut microbiome | 10.1002/imt2.20 | Yes | May 5, 2022 |
| From mechanism to application: decrypting light-regulated denitrifying microbiome through geometric deep learning | 10.21203/rs.3.rs-3086818/v1 | No | June, 2023 |
| MT-MAG: Accurate and interpretable machine learning based taxonomic assignment of metagenome-assembled genomes, with a partial classification option | 10.1101/2022.01.12.475159 | Yes | June 21, 2023 |
| A Virtual Machine Platform for Non-Computer Professionals for Using Deep Learning to Classify Biological Sequences of Metagenomic Data | 10.3791/62250 | No | September 25, 2021 |
| Deep Learning for Metagenomic Data: using 2D Embeddings and Convolutional Neural Networks | 10.48550/arXiv.1712.00244 | Yes | December, 2017 |
| Ontology-aware deep learning for antibiotic resistance gene prediction: novel function discovery and comprehensive profiling from metagenomic data | 10.1093/bib/bbac005 | Yes | March, 2022 |
| A Novel Metagenomic Binning Framework Using NLP Techniques in Feature Extraction | 10.2197/ipsjtbio.15.1 | Yes | January, 2022 |
| MEGGASENSE – The Metagenome/Genome Annotated Sequence Natural Language Search Engine: A Platform for the Construction of Sequence Data Warehouses | 10.17113/ftb.55.02.17.4749 | Yes | June, 2017 |
| Machine learning and deep learning applications in microbiome research | 10.1038/s43705-022-00182-9 | Yes | October 6, 2022 |
| Scalable learning of interpretable rules for the dynamic microbiome domain | 10.1101/2020.06.25.172270 | Yes | June 28, 2022 |
| Enhancing Microbiome Host Disease Prediction with Variational Autoencoders | 10.36837/chapman.000297 | Yes | July 1, 2021 |
| Marginalised stack denoising autoencoders for metagenomic data binning | 10.1109/CIBCB.2017.8058552 | Yes | August, 2017 |
| Human Gut Microbiome Data Analysis for Disease Likelihood Prediction Using Autoencoders | 10.1109/ICter53630.2021.9774811 | No | December, 2021 |
| MT-MAG: Accurate and interpretable machine learning for complete or partial taxonomic assignments of metagenome-assembled genomes | 10.1101/2022.01.12.475159 | Yes | January 12, 2022 |
| Deep Neural Network Modeling for Phenotypic Prediction of Metagenomic Samples | 10.1145/3388440.3414921 | Yes | November 10, 2022 |
| A Fast Word Embedding Based Classifier to Profile Target Gene Databases in Metagenomic Samples | 10.1007/978-3-030-79290-9_10 | No | July 3, 2021 |
| Reference-free biomarker mining in metagenomic data using language embedding | ad1c6a14-d8a0-4973-9eae-fac46b0bdf9c | Yes | April 21, 2021 |
| Brain neural network, development, microbiome, microbial toxins and COVID-19 | 10.52547/jcbior.3.1.14 | Yes | 2022 |
| Efficient and Interpretable Machine Learning Algorithms for Predictive Analyses in Metagenomic Data | 10.21203/rs.2.21634/v1 | Yes | January, 2020 |
| Scalable learning of interpretable rules for the dynamic microbiome domain | 10.1101/2020.06.25.172270 | Yes | June 28, 2020 |
| Enhanced Metagenomic Deep Learning for Disease Prediction and Reproducible Signature Identification by Restructured Microbiome 2D-Representations | 10.1016/j.patter.2022.100658 | No | January 13, 2023 |
| Taxonomic Profiling of Metagenomic Reads Using Deep Learning | 10.23860/thesis-Cres-Cecile-2020 | Yes | 2020 |
| A novel deep learning pipeline for early detection of colorectal cancer and colorectal adenoma using gut microbiome data | 10.1158/1538-7445.AM2023-3032 | No | April 4, 2023 |
| Deep learning for microbiome-based disease prediction and rheumatoid arthritis hand joint detection | 1993/37128 | No | December 15, 2022 |
| Efficient Sequence Clustering and Embedding Algorithms for Large-scale Metagenomics Data | 13884437 | Yes | 2019 |
| Some contributions to deep learning for metagenomcs | tel-01958389v1 | Yes | September 26, 2018 |
| Representation counts: the impact of embedding models on disease detection tasks from microbiome sequencing data | c0aaa51a-18b3-4e2a-b507-9cb4bd7c08a9 | Yes | June 23, 2022 |
| NLP-based classification of software tools for metagenomics sequencing data analysis into EDAM semantic annotation | 10.48550/arXiv.2210.00831 | Yes | October 18, 2022 |
| Multimodal deep learning models to classify healthy and disease states of human microbiome | 10.1038/s41598-022-04773-3 | Yes | January 17, 2022 |
| Advances in Microbiome Analysis: From the Variance Component Model to Deep Learning | 13886596 | Yes | 2019 |
| Deep learning-based colorectal cancer classification using augmented and normalised gut microbiome data | 14415 | Yes | 2022 |
| DeepBiome: a phylogenetic tree informed deep neural network for microbiome data analysis | 10.21203/rs.3.rs-33860/v1 | Yes | June 18, 2020 |
| Deep learning approach to metagenomic binning | 1721.1/119755 | Yes | 2018 |
| Deep Learning Frameworks for Multi-Omics Analyses of the Microbiome in Disease Studies | 19187285 | Yes | December 1, 2021 |
| Predicting the Complexity and Progression of the Gut Microbiome Using Temporal Data and Deep Learning | 8pt8g0bm | Yes | 2019 |
| Novel Mediation Method for Metagenomic Studies and Deep Learning Methods for Single Cell RNA-Seq Studies | 28258769 | Yes | 2020 |
| LA2SNE: A novel stochastic neighbor embedding approach for microbiome data visualization | 10.1109/BIBM.2014.6999294 | No | 2014 |
| A virtual machine platform for non-computer professionals for using deep learning to classify biological sequences of metagenomic data | 10.3791/62250 | No | September 25, 2021 |
| Disease Classification in Metagenomics with 2D Embeddings and Deep Learning | 10.48550/arXiv.1806.09046 | Yes | June 23, 2018 |
| Inferring Gut Microbial Interaction Network from Microbiome Data Using Network Embedding Algorithm | 10.12146/j.issn.2095-3135.20190704001 | Yes | October 9, 2019 |
| Constructing Long Short-Term Memory Networks to Predict Ulcerative Colitis Progression from Longitudinal Gut Microbiome Profiles | 10.33137/utjph.v2i2.36763 | Yes | September 5, 2021 |
| Female Urinary Metagenomic Analysis and Natural Language Processing Enhances the Infectious Diagnostic Yield in Precision Medicine | 10.1093/ofid/ofaa439.1859 | Yes | October 7, 2020 |
| META2: Memory-efficient taxonomic classification and abundance estimation for metagenomics with deep learning | 10.48550/arXiv.1909.13146 | Yes | February 10, 2020 |
| Deep Multiple Instance Learning for Taxonomic classification of metagenomic read sets | 10.48550/arXiv.1909.13146 | Yes | February 10, 2021 |
| Some contributions to deep learning for metagenomics | tel-01958389v1 | Yes | September 26, 2018 |
| Accurate identification of bacteriophages from metagenomic data using Transformer | 10.1093/bib/bbac258 | Yes | June 30, 2022 |
| ViBE: a hierarchical BERT model to identify eukaryotic viruses using metagenome sequencing data | 10.1093/bib/bbac204 | No | 2022 |
| A Deep Embedded Clustering Algorithm for the Binning of Metagenomic Sequences | 10.1109/ACCESS.2022.3176954 | Yes | May 23, 2022 |
| Virsearcher: Identifying Bacteriophages from Metagenomes by Combining Convolutional Neural Network and Gene Information | 10.1109/TCBB.2022.3161135 | No | 2023 |
| Phylogeny-Aware Deep 1-Dimensional Convolutional Neural Network for the Classification of Metagenomes | 10.1109/BIBM.2018.8621543 | No | January 24, 2019 |
| RNN-VirSeeker: A Deep Learning Method for Identification of Short Viral Sequences From Metagenomes | 10.1109/TCBB.2020.3044575 | Yes | June, 2022 |
| CLMB: deep contrastive learning for robust metagenomic binning | 10.48550/arXiv.2111.09656 | Yes | November 18, 2021 |
| Multi-Layer and Recursive Neural Networks for Metagenomic Classification | 10.1109/TNB.2015.2461219 | Yes | September 6, 2015 |
| Graph Embedding Deep Learning Guides Microbial Biomarkers' Identification | 10.3389/fgene.2019.01182 | Yes | November 22, 2019 |
| Enhancing Metagenome-based Disease Prediction by Unsupervised Binning Approaches | 10.1109/KSE.2019.8919295 | No | October, 2019 |
| MicroPheno: predicting environments and host phenotypes from 16S rRNA gene sequencing using a k-mer based representation of shallow sub-samples | 10.1093/bioinformatics/bty296 | Yes | June 27, 2018 |
| Diagnosis Approaches for Colorectal Cancer Using Manifold Learning and Deep Learning | 10.1007/s42979-020-00297-7 | No | August 27, 2020 |
| TaxoNN: ensemble of neural networks on stratified microbiome data for disease prediction | 10.1093/bioinformatics/btaa542 | Yes | May 25, 2020 |
| Multiclass Disease Classification from Microbial Whole-Community Metagenomes using Graph Convolutional Neural Networks | 10.1101/726901 | Yes | August 16, 2019 |
| Model-free prediction of microbiome compositions | 10.1101/2022.02.04.479107 | Yes | February 4, 2022 |
| DETIRE: a hybrid deep learning model for identifying viral sequences from metagenomes | 10.3389/fmicb.2023.1169791 | Yes | June 16, 2023 |
| A Framework for Effective Application of Machine Learning to Microbiome-Based Classification Problems | 10.1128/mbio.00434-20 | Yes | June 9, 2022 |
| Taxonomy-aware feature engineering for microbiome classification | 10.1186/s12859-018-2205-3 | Yes | June 15, 2018 |
| Resource saving taxonomy classification with k-mer distributions and machine learning | 10.48550/arXiv.2303.06154 | Yes | March 10, 2023 |
| Enhancing Disease Prediction on Imbalanced Metagenomic Dataset by Cost-Sensitive | 10.14569/IJACSA.2020.0110778 | Yes | 2020 |
| Applications of Machine Learning in Human Microbiome Studies: A Review on Feature Selection, Biomarker Identification, Disease Prediction and Treatment | 10.3389/fmicb.2021.634511 | Yes | February 19, 2021 |
| MITRE: predicting host status from microbiota time-series data | 10.1186/s13059-019-1788-y | Yes | September 2, 2019 |
| Efficient Discretization Approaches For Machine Learning Techniques To Improve Disease Classification On Gut Microbiome Composition Data | 10.25046/aj050368 | Yes | January, 2020 |
| The influence of machine learning technologies in gut microbiome research and cancer studies - A review | 10.1016/j.lfs.2022.121118 | Yes | December 15, 2022 |
| VirNet: Deep attention model for viral reads identification | 10.1109/ICCES.2018.8639400 | No | December, 2018 |
| The Phylogenetic Tree based Deep Forest for Metagenomic Data Classification | 10.1109/BIBM.2018.8621463 | No | December, 2018 |
| An Ensemble Feature Selection Method Based on Deep Forest for Microbiome-Wide Association Studies | 10.1109/BIBM.2018.8621461 | No | January, 2019 |
| Gene Family Abundance Visualization based on Feature Selection Combined Deep Learning to Improve Disease Diagnosis | 10.5614/j.eng.technol.sci.2021.53.1.9 | No | January 30, 2021 |
| Effective Disease Prediction on Gene Family Abundance Using Feature Selection and Binning Approach | 10.1007/978-981-15-9354-3_2 | No | December 8, 2020 |

Table S 14: **Table of data availability.** This table contains the studies synthesized in this work, along with their date of publication, their id and whether they are publicly available or not.

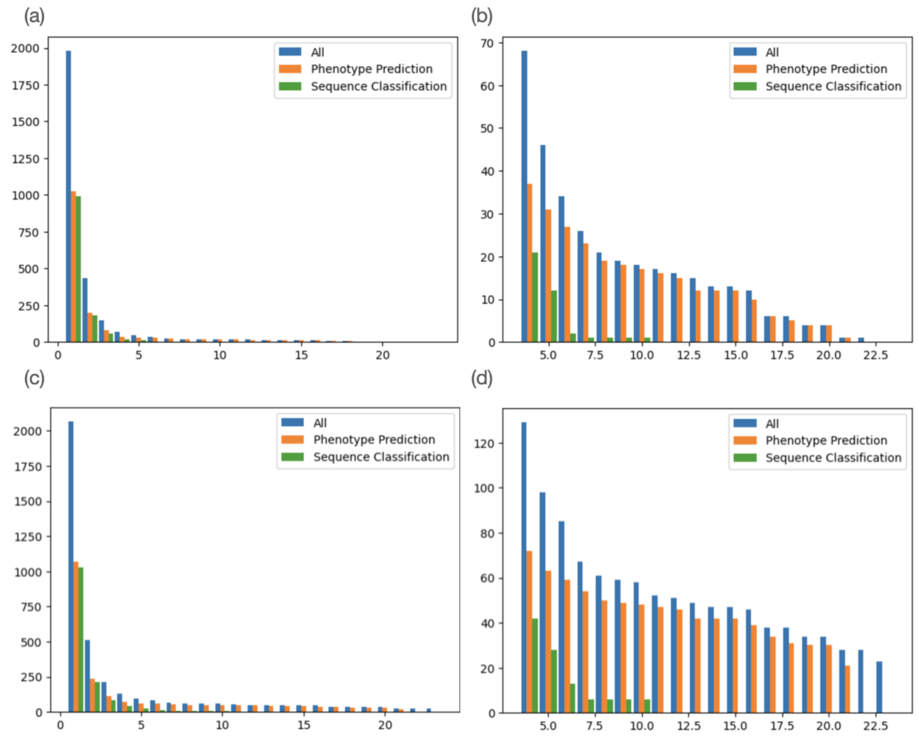


Fig S 2: This figure shows the number of articles according to the number of links from original articles pointing to them. ”Sequence classification” values mean the number of articles pointed by one of the articles of this category, same for ”Phenotype prediction”. ”All” represents all articles pointed by those two categories + 4 miscellaneous articles. **(a)** represents the distribution of articles through the number of times they are pointed at, **(b)** is a zoom of the precedent figure, starting with a threshold of minimum 4 citations. **(c)** and **(d)** represent the same thing but considering only newly discovered articles and not the ones already present in the database.

were more connected than the ones who did not pass this check. We plotted the proportion for each co-citation connectivity in our three datasets, the result can be seen in **Figure S4**. Although the number of retained articles is quite low, it seems that co-citation connectivity did not influence the relevance of an article. The distributions have close means (8 and 8.41 for ”All” and ”All After Filtering (AF)”, 10.68 and 10 for ”Phenotype Prediction” and ”Phenotype Prediction AF”,

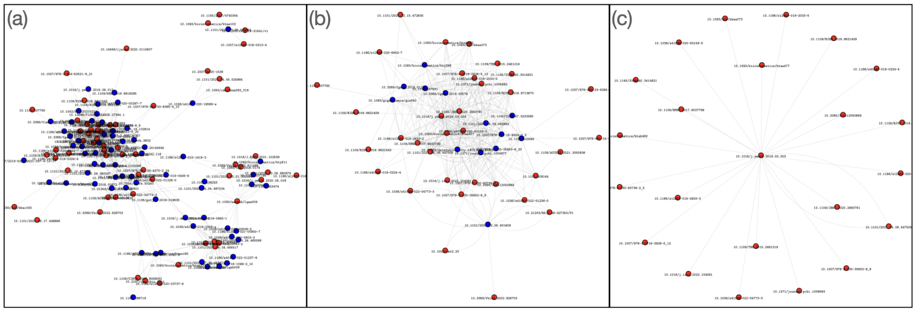


Fig S 3: Examples of generated graphs of articles : a vertex is an article (labelled by DOI). Red dots represent articles that were already selected by research equation, blue dots represent articles discovered through Connected Papers. Only articles pointed by a certain minimal number of links defined before. These articles consider the total of all articles, not only the "Sequence classification" or "Phenotype prediction" selections. (a) represents articles pointed by at least 4 links, (b) articles pointed by at least 14 articles and (c) articles pointed by at least 23 articles. We can see in the latter that only already presented articles remain, showing our research equation already had a firm grasp on the core of our subject

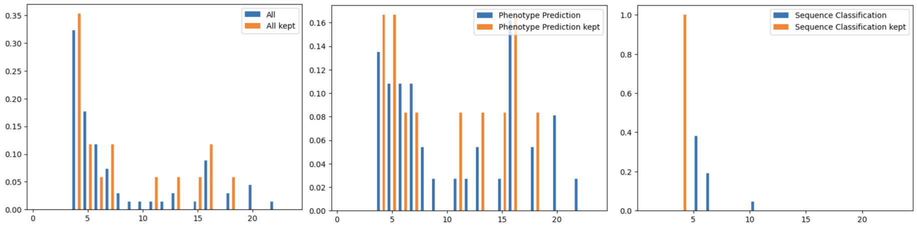


Fig S 4: The proportion of each connectivity in newly discovered articles and articles that passed the last filter

and finally 5.05 and 4 for "Sequence Classification" and "Sequence Classification AF"). We may note that most of the cited articles did not pass the filtering. This could be due to the fact that many cited articles tend to be more general and therefore may not be related to both domains. There is an environment of articles that are found many times but actually do not link both our topics

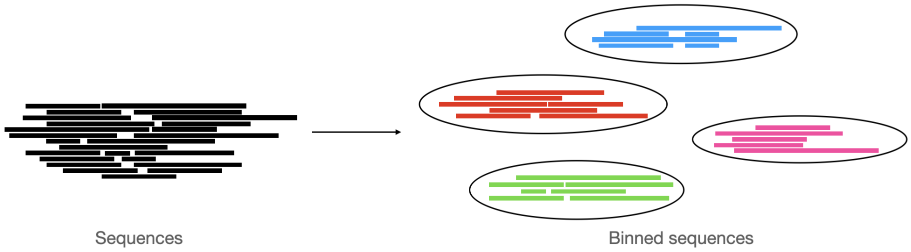


Fig S 5: Binning : grouping of sequences in different bins based on similarity

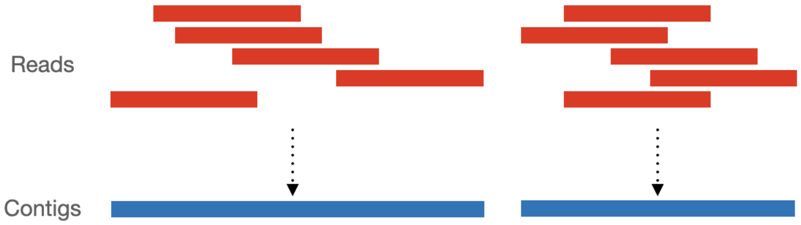


Fig S 6: Overlapping reads are combined into contigs

[2] Busia A, Dahl GE, Fannjiang C, Alexander DH, Dorfman E, Poplin R, et al. A deep learning approach to pattern recognition for short DNA sequences; p. 12.

[3] Matougui B, Batouche M, Boukelia A. A K-mer based Multi Convolutional Neural Network Classifier of Low-Ranking Taxonomic Bins from Metagenome; p. 13.

[4] Wijegunarathna K, Maduranga U, Weerasinghe S, Perera I, Wickaramarachchi A. Cluster Identification in Metagenomics – A Novel Tech-

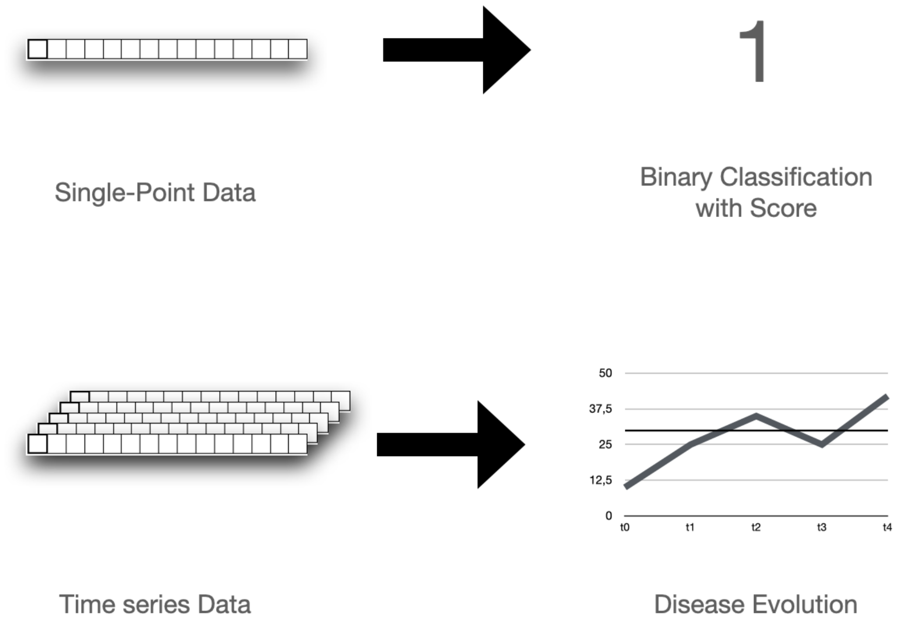


Fig S 7: Difference of paradigm between single-point analysis and longitudinal analysis

nique of Dimensionality Reduction through Autoencoders. International Journal on Advances in ICT for Emerging Regions (ICTer). 2021;14(2):9. doi:10.4038/icter.v14i2.7224.

[5] Menegaux R, Vert JP. Continuous Embeddings of DNA Sequencing Reads and Application to Metagenomics. Journal of Computational Biology. 2019;26(6):509–518. doi:10.1089/cmb.2018.0174.

[6] Fiannaca A, La Paglia L, La Rosa M, Lo Bosco G, Renda G, Rizzo R, et al. Deep learning models for bacteria taxonomic classification of metagenomic data. BMC Bioinformatics. 2018;19(S7):198. doi:10.1186/s12859-018-2182-6.

[7] Liang Q, Bible PW, Liu Y, Zou B, Wei L. DeepMicrobes: taxonomic classification for metagenomics with deep learning. NAR Genomics and

Bioinformatics. 2020;2(1):lqaa009. doi:10.1093/nargab/lqaa009.

[8] Maduranga U, Wijegunarathna K, Weerasinghe S, Perera I, Wickramarachchi A. Dimensionality Reduction for Cluster Identification in Metagenomics using Autoencoders. In: 2020 20th International Conference on Advances in ICT for Emerging Regions (ICTer). Colombo, Sri Lanka: IEEE; 2020. p. 113–118. Available from: https://ieeexplore.ieee.org/document/9325447/.

[9] Menegaux R, Vert JP. Embedding the de Bruijn graph, and applications to metagenomics. Bioinformatics; 2020. Available from: http://biorxiv.org/lookup/doi/10.1101/2020.03.06.980979.

[10] Rojas-Carulla M, Tolstikhin I, Luque G, Youngblut N, Ley R, Schölkopf B. GeNet: Deep Representations for Metagenomics; p. 13.

[11] Kouchaki S, Tirunagari S, Tapinos A, Robertson DL. Marginalised stack denoising autoencoders for metagenomic data binning. In: 2017 IEEE Conference on Computational Intelligence in Bioinformatics and Computational Biology (CIBCB). Manchester, United Kingdom: IEEE; 2017. p. 1–6. Available from: http://ieeexplore.ieee.org/document/8058552/.

[12] Georgiou A, Fortuin V, Mustafa H, Rätsch G. META\$ˆ\mathbf{2}\$: Memory-efficient taxonomic classification and abundance estimation for metagenomics with deep learning; 2020. Available from: http://arxiv.org/abs/1909.13146.

[13] Liang Kc. MetaVelvet-DL: a MetaVelvet deep learning extension for de novo metagenome assembly. 2021; p. 21.

[14] Matougui B, Boukelia A, Belhadef H, Galiez C, Batouche M. NLP-MeTaxa: A Natural Language Processing Approach for Metagenomic Taxonomic

Binning Based on Deep Learning. Current Bioinformatics. 2021;16(7):992–1003. doi:10.2174/1574893616666210621101150.

[15] Mock F, Kretschmer F, Kriese A, Böcker S, Marz M. BERTax: taxonomic classification of DNA sequences with Deep Neural Networks. Bioinformatics; 2021. Available from: http://biorxiv.org/lookup/doi/10.1101/2021.07.09.451778.

[16] Karagöz MA, Nalbantoglu OU. Taxonomic classification of metagenomic sequences from Relative Abundance Index profiles using deep learning. Biomedical Signal Processing and Control. 2021;67:102539. doi:10.1016/j.bspc.2021.102539.

[17] Nissen JN, Johansen J, Allesøe RL, Sønderby CK, Armenteros JJA, Grønbech CH, et al. Improved metagenome binning and assembly using deep variational autoencoders. Nature Biotechnology. 2021;39(5):555–560. doi:10.1038/s41587-020-00777-4.

[18] Zhang P, Jiang Z, Wang Y, Li Y. CLMB: deep contrastive learning for robust metagenomic binning; p. 20.

[19] Pan S, Zhu C, Zhao XM, Coelho LP. A deep siamese neural network improves metagenome-assembled genomes in microbiome datasets across different environments. Nature Communications. 2022;13(1):2326. doi:10.1038/s41467-022-29843-y.

[20] Kang DD, Li F, Kirton E, Thomas A, Egan R, An H, et al. MetaBAT 2: an adaptive binning algorithm for robust and efficient genome reconstruction from metagenome assemblies. PeerJ. 2019;7:e7359. doi:10.7717/peerj.7359.

[21] Piera Lindez P, Johansen J, Sigurdsson AI, Nissen JN, Rasmussen S. Adversarial and variational autoencoders improve metagenomic binning. Bioin-

formatics; 2023. Available from: http://biorxiv.org/lookup/doi/10.1101/2023.02.27.527078.

[22] Lamurias A, Tibo A, Hose K, Albertsen M, Nielsen TD. Metagenomic Binning using Connectivity-constrained Variational Autoencoders;.

[31] Rahman MA, Rangwala H. IDMIL: an alignment-free Interpretable Deep Multiple Instance Learning (MIL) for predicting disease from whole-metagenomic data; p. 9.

[32] Queyrel M, Prifti E, Templier A, Zucker JD. Towards end-to-end disease prediction from raw metagenomic data. Genomics; 2020. Available from: http://biorxiv.org/lookup/doi/10.1101/2020.10.29.360297.

[33] Strocchi M, Corso G, Liò P. Representation counts: the impact of embedding models on disease detection tasks from microbiome sequencing data; p. 12.

[42] Shen WX, Liang SR, Jiang YY, Chen YZ. Enhanced metagenomic deep learning for disease prediction and consistent signature recognition by

restructured microbiome 2D representations. Patterns. 2023;4(1):100658. doi:10.1016/j.patter.2022.100658.

[43] Nguyen TH, Prifti E, Chevaleyre Y, Sokolovska N, Zucker JD. Disease Classification in Metagenomics with 2D Embeddings and Deep Learning. arXiv:180609046 [cs]. 2018;.

[44] Nguyen HT, Bao T, Hoang H, Phuoc T, C N. Improving Disease Prediction using Shallow Convolutional Neural Networks on Metagenomic Data Visualizations based on Mean-Shift Clustering Algorithm. International Journal of Advanced Computer Science and Applications. 2020;11(6). doi:10.14569/IJACSA.2020.0110607.

[45] Pfeil J, Siptroth J, Pospisil H, Frohme M, Hufert FT, Moskalenko O, et al. Classification of Microbiome Data from Type 2 Diabetes Mellitus Individuals with Deep Learning Image Recognition. Big Data and Cognitive Computing. 2023;7(1):51. doi:10.3390/bdcc7010051.

[46] Mreyoud Y, Song M, Lim J, Ahn TH. MegaD: Deep Learning for Rapid and Accurate Disease Status Prediction of Metagenomic Samples. Life. 2022;12(5):669. doi:10.3390/life12050669.

[47] Fung DLX, Li X, Leung CK, Hu P. A self-knowledge distillation-driven CNN-LSTM model for predicting disease outcomes using longitudinal microbiome data. Bioinformatics Advances. 2023;3(1):vbad059. doi:10.1093/bioadv/vbad059.
